# Three amphioxus reference genomes reveal gene and chromosome evolution of chordates

**DOI:** 10.1101/2022.01.04.475009

**Authors:** Zhen Huang, Luohao Xu, Cheng Cai, Yitao Zhou, Jing Liu, Zexian Zhu, Wen Kang, Duo Chen, Surui Pei, Ting Xue, Wan Cen, Chenggang Shi, Xiaotong Wu, Yongji Huang, Chaohua Xu, Yanan Yan, Ying Yang, Wenjing He, Xuefeng Hu, Yanding Zhang, Youqiang Chen, Changwei Bi, Chunpeng He, Lingzhan Xue, Shijun Xiao, Zhicao Yue, Yu Jiang, Jr-Kai Yu, Erich D. Jarvis, Guang Li, Gang Lin, Qiujin Zhang, Qi Zhou

## Abstract

The slow-evolving invertebrate amphioxus has an irreplaceable role in advancing our understanding into the vertebrate origin and innovations. Here we resolve the nearly complete chromosomal genomes of three amphioxus species, one of which best recapitulates the 17 chordate ancestor linkage groups. We reconstruct the fusions, retention or rearrangements between descendants of whole genome duplications (WGDs), which gave rise to the extant microchromosomes likely existed in the vertebrate ancestor. Similar to vertebrates, the amphioxus genome gradually establishes its 3D chromatin architecture at the onset of zygotic activation, and forms two topologically associated domains at the *Hox* gene cluster. We find that all three amphioxus species have ZW sex chromosomes with little sequence differentiation, and their putative sex-determining regions are nonhomologous to each other. Our results illuminate the unappreciated interspecific diversity and developmental dynamics of amphioxus genomes, and provide high-quality references for understanding the mechanisms of chordate functional genome evolution.

## Introduction

Although first described in 1774, the lesser-known marine invertebrate amphioxus (or lancelets) only became recognized for its unparalleled value in elucidating the vertebrate origin and innovations until characterization of its *Hox* genes in 1990s (*1*). It was later established that amphioxus diverged from the ancestor of two other chordate subphyla, urochordates (tunicates) and vertebrates about 550 million years ago (MYA) (*2, 3*). Amphioxus has a vertebrate-like but simpler body plan, and underwent much less lineage-specific changes of chromosomes and genomic sequences than urochordates (*4*). Therefore, it represents the best known living proxy for the chordate ancestor (*5, 6*). Amphioxus has one, and the largest reported *Hox* gene cluster with 15 genes (*7*), which was found to form one structural and regulatory unit of topologically associated domain (TAD). By contrast, vertebrates have at least 4 *Hox* gene clusters and up to 13 genes per cluster, with the mouse *HoxA* and *HoxD* clusters each forming two TADs (*8*). Such a 4-fold difference of *Hox* gene cluster numbers provided early evidence for Ohno’s hypothesis of two rounds of WGDs (the 2R hypothesis) (*9, 10*) that shaped the genome evolution and regulation of vertebrates since they diverged from other chordates.

Broader understanding beyond individual genes into the scenario and functional consequences of vertebrate WGDs, whose times and timing recently became a subject of debate (*11*), necessitate high-quality sequence assembly and annotation of genes and *cis*-regulatory elements of amphioxus (*12*), as a pre-WGD outgroup. The first draft genome of Florida amphioxus *Branchiostoma floridae* (Bf) was published over a decade ago, and has been frequently used to reconstruct the ancestral vertebrate protokaryotype, with however different estimates of ancestral linkage group number between studies (*11, 13–15*). A recent work improved the Bf genome into the chromosome-level and proposed a refined 2R hypothesis with 17 ancestral chordate linkage groups: the first WGD occurred in the ancestor of all vertebrates, and the second WGD only occurred in the lineage of jawed vertebrates (*16*). The duplicated gene products of WGDs in vertebrates (‘ohnologues’) seem to have generally a higher number of and more specialized regulatory elements and gene expression between copies, relative to their single-copy orthologs of amphioxus (*12*). Besides results at the gene-level, to address how vertebrates evolved globally more complex regulatory circuits after WGDs requires knowledge of higher-order chromatin organization of amphioxus.

An often-overlooked factor among previous studies using only one species’ genome is amphioxus’ largely unexplored interspecific genomic diversity. It is known that different amphioxus species have different chromosome numbers, and exhibit frequent disruptions of gene synteny which may confound the inference of vertebrate ancestral state (*17*). Moreover, the available amphioxus genome assemblies are either incomplete or fragmented because of the high intraspecific polymorphisms associated with their large effective population size (*4*). To elucidate the evolution of genes, genomes and chromatin landscapes of different amphioxus species compared to vertebrates, we resolved here the nearly complete haploid genomes of three *Branchiostoma* amphioxus species Chinese amphioxus (*B. belcheri*, Bb), Japanese amphioxus (*B. japonicum*, Bj) and Bf.

## Results

### Haploid chromosomal genomes of three amphioxus species

We estimated the genome-wide heterozygosity levels of three amphioxus species to range from 3.2% to 4.2%, among the highest in animal species (*18*) (**Supplementary Fig. S1**). To overcome this great challenge for genome assembly, we devised an interspecific trio-sequencing strategy and produced respectively more than 100-fold short and long sequencing reads for the F1 hybrids derived from Bf-Bb or Bf-Bj crosses (**Fig. 1a**, **Supplementary Fig. S2**). Given at least 50 MYs’ species divergence time (**Supplementary Fig. S3**), the hybrids contain two haploid parental genomes that have become too diverged in sequences to form cross-species chimeric assembly (**Supplementary Fig. S1**). By mapping short-reads derived from the respective parental species, we were able to attribute each assembled contig into one of the four haploid (Bb, Bj and two Bf) genomes (**Fig. 1b-c**). The new haploid amphioxus genomes have an assembled size ranging from 382 to 491 Mb, and an over 200-fold improvement in contig N50 length (between 6.4 to 14.2Mb) compared to the published genomes(*4, 12, 17*), an over 97% genome completeness (measured by BUSCO) and a reduced level of false duplications (**Supplementary Table S1, Supplementary Fig. S4a-b**). Using Hi-C data, we anchored more than 98.6% of the contig sequences into chromosomes, with a much lower gap number (on average only 3.8 gaps) per chromosome than those of major vertebrate reference genomes and that of a recently improved Bf genome(*16*) (**Fig. 1d**, **Supplementary Fig. S4c**).

**Figure 1.**
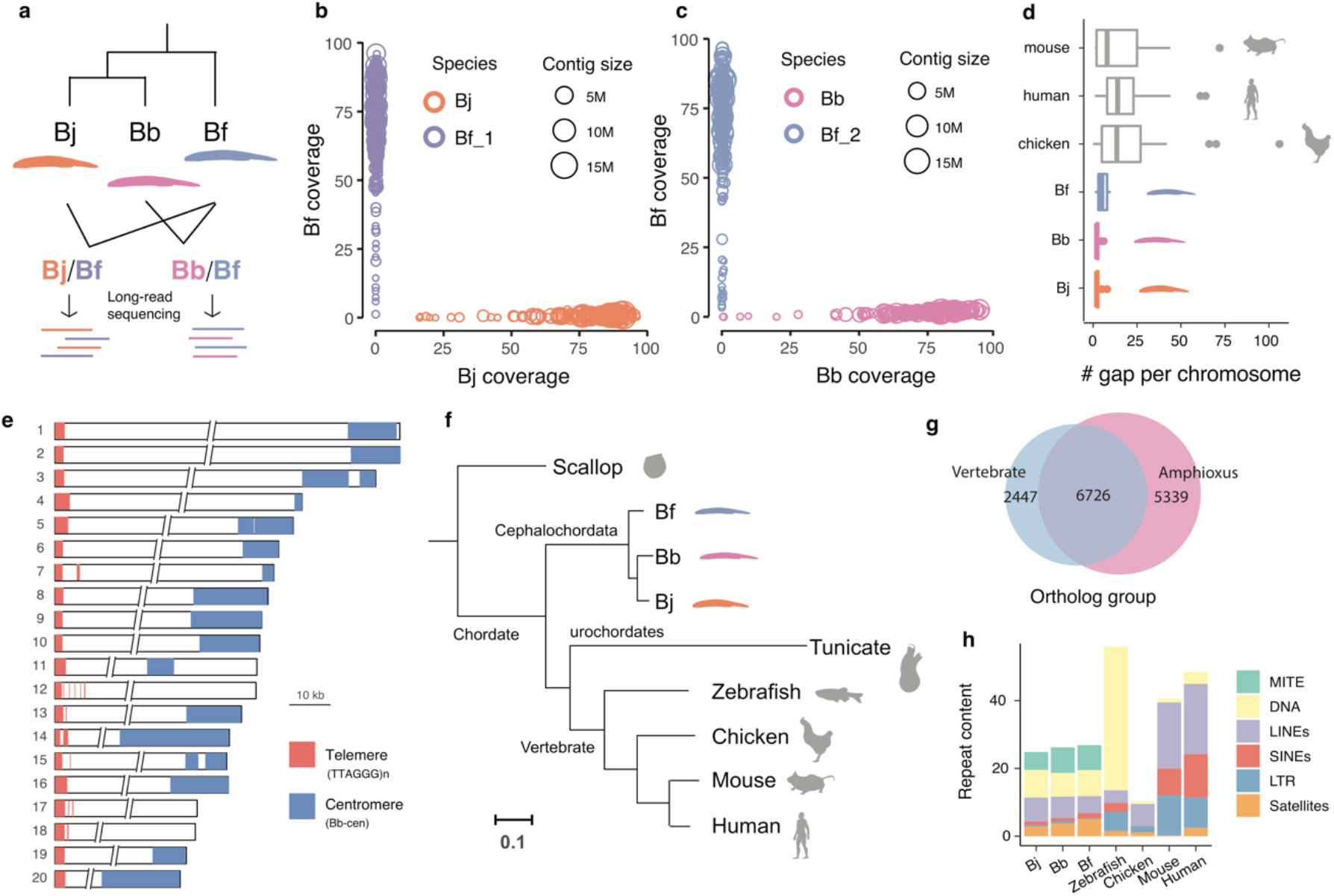
Three haploid genomes of amphioxus species. a) We performed long-read sequencing of interspecific hybrids between the three amphioxus species and assembled their haploid genomes. Bj: *B. japonicum*, Bb: *B. belcheri*, Bf: *B. floridae*. **b-c)** Contig sequences of the hybrids were assigned to the haploid genome of each parental species, according to their coverage (proportion of mapped sequences) mapped by the short-reads of parental species. **d)** The amphioxus haploid genomes have a lower gap content (numbers of gaps per chromosome) compared to other vertebrate reference genomes. **e)** Most amphioxus chromosomes are telocentric. The 10kb scale applies to the two tips of the chromosomes only, and the two slash lines represent the gaps between the two chromosomal tips. **f)** Phylogenomic tree based on whole-genome alignments of amphioxus vs. other chordate species. **g)** A large number of orthologous gene groups (6726) is shared between amphioxus and vertebrates, but amphioxus species have 5,339 specific gene groups. **h)** MITEs (green) comprise ∼6.7% of the amphioxus genomes but are largely absent in vertebrates. In the DNA transposon category (yellow) MITE was excluded.

With some exceptions, all chromosomal sequences of the three species have been assembled from the telomere at one end to the centromere at the other (**Fig. 1e, Supplementary Fig. S5-6**). This is consistent with the reported predominantly telocentric karyotype of amphioxus (*19–21*), the low levels of recombination rate and nucleotide diversity at centromeric and pericentromeric regions (**Supplementary Fig. S7-8**); and is also verified by our fluorescent *in situ* hybridization (FISH) experiment for Bf (**Supplementary Fig. S9**). The telomeres contain conserved telomeric motifs (TTAGGG)_n_ (*22*) with an average length of 3.6 kb, and they account for the majority of G-quadruplex content in the genome (**Supplementary Fig. S10**). Our cytogenetic and genomic investigations also confirmed the presence of interstitial telomeric sequences in a few amphioxus chromosomes (**Supplementary Fig. S5, S9**). The putative centromeric regions consist of species-specific satellite monomers of different sequences and lengths, with inverted repeat structures (**Supplementary Fig. S11**). Besides the telomeres and centromeres, our new genomes contain newly resolved complex repeat regions, including satellite DNA or rDNA arrays (**Supplementary Fig. S12**), that are partial or absent in the previous amphioxus genomes.

Our phylogenomic analyses using whole-genome alignments of amphioxus against other chordates and one invertebrate outgroup confirmed amphioxus as the most basal chordate lineage, with a relatively lower genome-wide substitution rate (**Fig. 1f**). Based on 3,653 single-copy orthologous genes, we estimated that different chordate lineages diverged about 552.0 MYA, and three amphioxus species diverged about 86.6 MYA (**Supplementary Fig. S3**). Over 73% vertebrate orthologous gene groups are present in amphioxus genomes (**Fig. 1g**). The vertebrate specific genes are enriched for various gene ontology (GO) categories including signalling pathway regulation and muscle functions, while the amphioxus specific genes are enriched for GOs of tissue regeneration (*23*) and apoptosis, among many others (**Supplementary Table S2**). We also identified 27,032 conserved sequence elements between vertebrates and amphioxus, and majorities of them (26,955) are located in protein-coding regions. Finally, the amphioxus genomes were found to have a moderate repeat content of about 30% (**Fig. 1h**), but they contain abundant MITEs (miniature inverted-repeat transposable elements) that are nearly absent in vertebrates. These MITEs seem to have propagated more recently in amphioxus species, relative to other DNA transposons (**Supplementary Fig. S13**).

### Ancestral karyotypes of amphioxus, chordates and vertebrates

The assembled chromosome number of Bj, Bf and Bb is respectively 18, 19 and 20, consistent with their reported karyotypes (*22, 24*). Based on their whole-genome alignments, we inferred that similar to the karyotype of Bb, the *Branchiostoma* amphioxus ancestor had 20 linkage groups, which then underwent two chromosome fusions in Bj, and one fusion in Bf after their species divergence (**Fig. 2a, Supplementary Fig. S14**).

**Figure 2.**
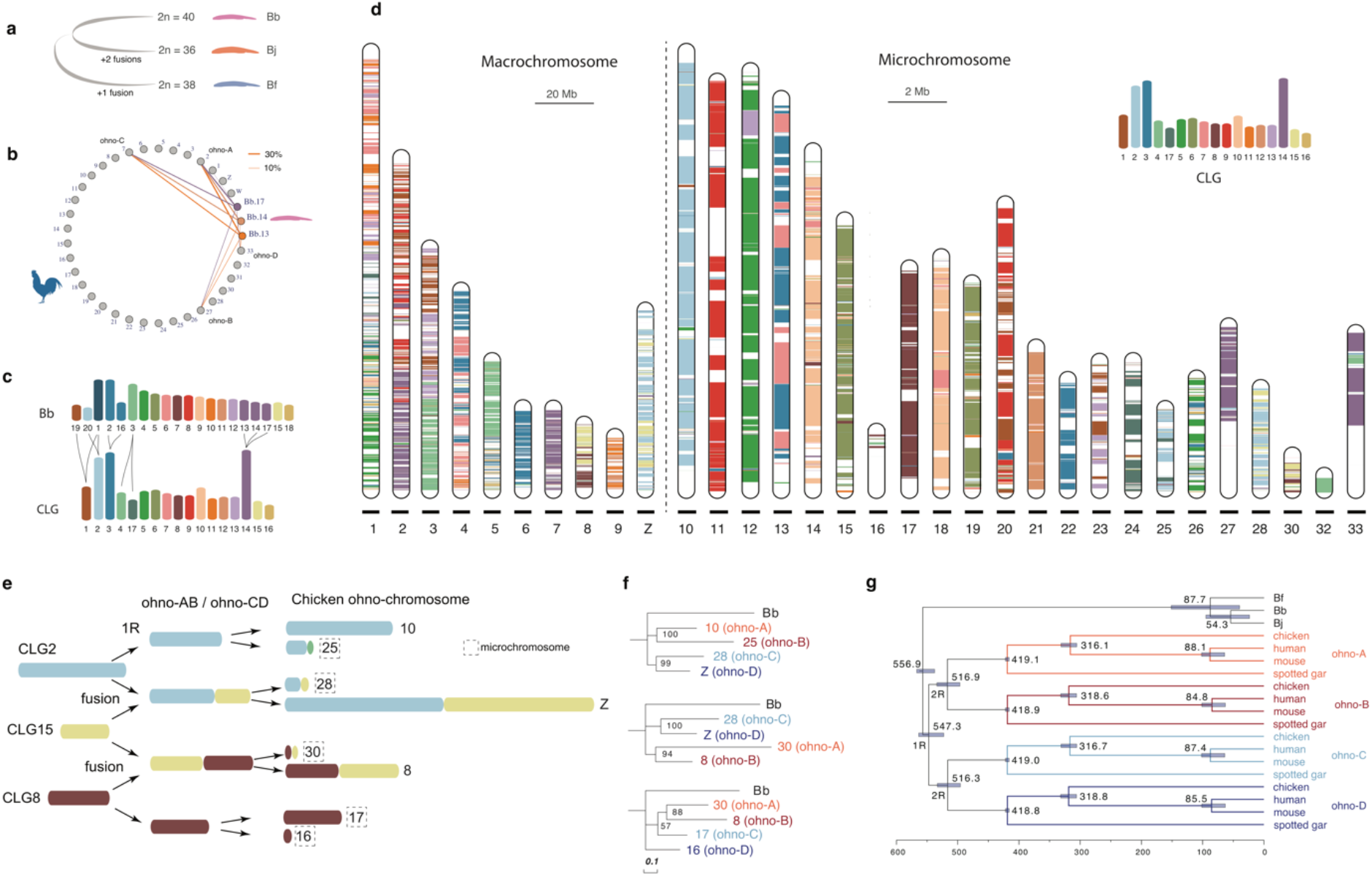
Ancestral karyotypes of amphioxus, chordates and vertebrates. a) Bb probably best recapitulates the ancestral karyotype of *Branchiostoma* amphioxus, with Bj and Bf having undergone chromosomal fusions. **b)** Genes on chr13, chr14 and chr17 of Bb have their homologous genes located on the same set of chicken chromosomes. Each line connecting chromosomes of Bb and chicken chromosomes is scaled to the proportion of Bb genes that are homologous to the genes of one chicken chromosome. **c)** The inferred relationship between Bb chromosomes and CLG. **d)** Composition of chicken chromosome by CLG homologous sequences. The colored bands represent the Bb-chicken synteny blocks. A different scale for macrochromosomes (20 Mb) and microchromosomes (2 Mb) was used. **e)** Reconstructed 1R and 2R of three CLGs. One color represents one CLG, and when one chromosome is composed with more than one CLG, two or more CLG blocks are linked together. **f)** The ohnolog genes were used to construct the phylogeny of ohno-chromosomes (ohno-A, B, C, D), which refer to gene groups derived from WGDs. Bb homologs were used as the outgroup. Bootstrapping values shown placed at the internal nodes. **g)** 244 ohnolog gene groups were used to date 1R and 2R. Fossil calibration for the mouse-human nodes: 62-101 MY, bird-mammal nodes: 306-332 MY.

Genomic comparison between Bb vs. chicken allows us to reconstruct the karyotype of chordate ancestor. We chose chicken because it is one of the vertebrates that exhibit the lowest rates of lineage-specific chromosomal evolution (*15, 16, 25, 26*) and gene duplications (*27, 28*). Consistent with two rounds of WGDs followed by gene loss, one single-copy amphioxus gene typically has between one to four homologs in vertebrates (**Supplementary Fig. S15**). Moreover, genes from one Bb chromosome are more frequently found to have homologs distributed on four different chromosomes in chicken (**Supplementary Fig. S16**), compared to spotted gar or human (**Supplementary Fig. S17**), confirming that chicken better preserve the ancestral vertebrate karyotype with less interchromosomal rearrangements. We also found several Bb chromosomes share their combination of homologous chicken chromosomes. For instance, Bb chr13, chr14 and chr17 all have their homologous genes located on the chicken chr2 (GGA2), GGA7, GGA27 and GGA33 (**Fig. 2b**). This suggested that these three Bb chromosomes were likely derived from one single chordate ancestral linkage group (CLG) (**Fig. 2c**). Similarly, Bb chr1 shares its homologous chicken chromosomes exclusively with either Bb chr19 or chr20 (**Supplementary Fig. S16,** **Fig. 2c**), suggesting Bb chr1 originated from a translocation between two CLGs. Moreover, we inferred that Bb chr2 and chr16 fused at the vertebrate ancestor prior to whole genome duplication, while Bb chr3 was split into two (**Supplementary Fig. S16,** **Fig. 2c**). Taken together, we inferred that there was a total of 17 CLGs (**Fig. 2c**), consistent with previous results (*4, 13, 16*).

To reconstruct the evolutionary trajectories of how CLGs gave rise to the representative extant vertebrate karyotypes, we mapped the homologs of Bb genes assigned to 17 CLGs (**Fig. 2c**) across the chromosomes of chicken or spotted gar. Most chicken and gar microchromosomes have homologous Bb genes predominantly derived from one single CLG (**Fig. 2d, Supplementary Fig. S18**). Such striking evolutionary stability of microchromosomes spanning the entire chordate evolution supports the hypothesis that they were likely present at the ancestor of bony vertebrates (*16, 29–31*). Some chicken microchromosomes (e.g., GGA28 and GGA30), like most macrochromosomes, nevertheless are homologous to two or more CLGs (**Fig. 2d**). When the same combination of CLGs were found for two different homologous GGAs, e.g., GGA28 and GGAZ (homologous to CLG2 and CLG15), we inferred a fusion or translocation likely occurred between 1R and 2R, as illustrated in **Fig. 2e**. We identified a total of 5 such putative post-1R chromosome fusions or translocations (**Supplementary Fig. S19**), whose 2R descendant genes are predicted to be grouped together (**Fig. 2f**, e.g., GGA28 and GGAZ genes) apart from other ohnologs (GGA10 and GGA25 genes) of the same CLG origin but without undergoing post-1R fusions or translocations. This was broadly supported by the phylogenetic trees (**Fig. 2f**, **Supplementary Fig. S19)** constructed from chicken ohnolog gene groups (**Supplementary Table S3, Supplementary Fig. S20**). Extending our phylogenetic reconstructions to 243 chicken paralog groups with at least three ohnologs available, we found among the 9 CLGs that gave rise to ohnologs distributed on 4 GGAs (we termed genes of each of these 4 GGAs as ‘ohno linkage group’, ohno-A, B, C, D), 7 CLGs’ ohnolog trees exhibited a phylogenetic structure that strongly supported the 2R hypothesis (**Supplementary Fig. S21**). That is, ohnologs from two GGAs of the same post-1R origin (ohno-A/B or C/D) were grouped together in their phylogenetic trees. When such ohno linkage groups involve microchromosomes, we revealed that microchromosomes always contain much less ohnologs than the other macrochromosomes of the same post-1R origin (**Supplementary Fig. S22**). This led to our hypothesis that microchromosomes possibly originated by asymmetric sequence loss after the 2R in the vertebrate ancestor.

By concatenating chicken ohnologs from the same ohno linkage group (A, B, C or D), together with their orthologs of human, mouse and gar, we constructed their phylogenetic trees and dated the timing of 1R and 2R (**Fig. 2g**). The 1R was estimated to occur 547 MY ago, in less than 10 MY since the divergence of chordate common ancestor (**Fig. 2g**). In addition, we estimated that jawed vertebrates experienced 2R about 517 MY ago (**Fig. 2g**), 10 MY after their divergence from jawless vertebrates (**Supplementary Fig. S3**).

### Amphioxus specific gene duplications

Although without undergoing WGDs, the three amphioxus species have a comparable number of protein-coding genes (between 22,733 to 26,497) to that of vertebrates (**Supplementary Fig. S23a**). By phylogenetic reconstruction of 8,464 orthologous gene groups whose members are present in both amphioxus and vertebrates, we estimated that the amphioxus ancestor had acquired 4,855 genes (**Fig. 3a**), some of which may also result from gene loss in the vertebrate ancestor. Interestingly, genes that retained at least two paralogs in vertebrates are more likely to have undergone duplications in amphioxus (*P* < 1.71e-13, Fisher’s exact test, **Supplementary Table S4**), suggesting convergent gene gains in vertebrates and amphioxus. For example, among the orthologous gene groups that have multi-copy genes in Bb, 74% have multi-copy homologs in chicken, but only 33% of the orthologous gene groups with single-copy Bb genes have homologs in chicken (**Fig. 3b**). We also found cases of recurrent duplication in amphioxus (**Supplementary Fig. S23b**) as demonstrated by a recent study for *MRF* genes (*32*). For instance, there are 3 ohnologs of the *Slc27a* gene family derived from a single chordate ancestral gene which was independently duplicated multiple times at the ancestor of amphioxus (**Fig. 3c**).

**Figure 3.**
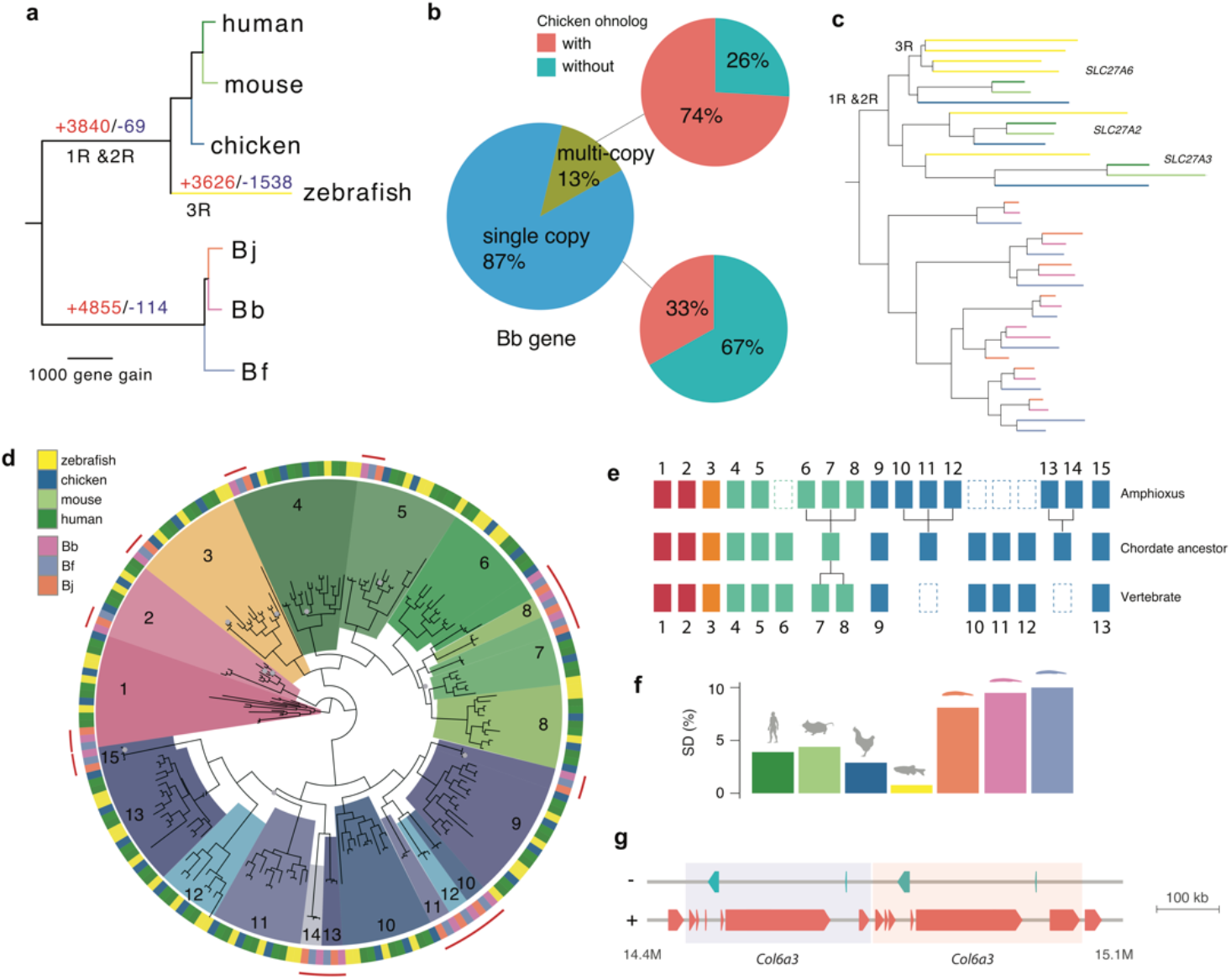
Gene expansion in amphioxus species a) Reconstructed gene gains (red) and losses (blue) events during the chordate evolution based on the ortholog gene groups. The branch length is scaled to the number of gene gain. **b)** Using Bb as an example, we show duplicated genes in amphioxus more frequently have paralogs in vertebrates. A majority (74%) of the Bb multi-copy genes have chicken paralogs compared with only 33% of Bb single-copy genes. **c)** Independent expansion of *SLC27A* gene copies in vertebrates (due to WGD) and amphioxus (due to gene duplication). Each species (zebrafish and three amphioxus species) is marked with the same color as shown in a). **d)** Phylogenetic tree of *Hox* genes. The homologous *Hox* gene (denoted by the number) group of amphioxus and vertebrates was marked in the same color. The grey dots at the internal nodes indicate a bootstrapping value lower than 60. **e)** An inferred model of *Hox* gene evolution in chordates according to the results of d). Dashed boxes denote gene loss, each aligned column denotes homologous relationship, individual gene duplications are also shown for either amphioxus or vertebrates. **f)** Amphioxus has a higher portion of genome derived from segmental duplication compared to vertebrates **g)** One example of segmental duplication involving *Col6a3* in Bf. The two copies are next to each other highlighted in different background colors

The other prominent case of convergent gene acquisition in amphioxus and vertebrates is demonstrated by certain members of *Hox* genes. Amphioxus has one prototypical *Hox* gene cluster (*AmphiHox*), whose posterior *Hox* genes have an ambiguous orthologous relationship with the vertebrate *Hox* paralog groups (HPGs), leaving the *Hox* gene number of chordate ancestor still controversial (*33–35*). Our phylogenetic analysis confirmed one-to-one homologous relationships of some *Hox* (1-5, 9, 15) genes between amphioxus and vertebrates, dating their likely existence to the chordate ancestor (**Fig. 3d**). Other *Hox* genes likely have undergone gain and loss events independently in the ancestors of the two clades’ (**Fig. 3e**). For instance, the amphioxus *Hox6-8* and the vertebrate *HPG8* seem to be acquired after the two chordate clades diverged from each other. The posterior amphioxus *Hox* genes *Hox10-12* and *Hox13-14* are respectively grouped with the vertebrate *HPG9* and *HPG11-13*, suggesting amphioxus-specific duplications from an ancestral chordate *Hox* gene that might have subsequently become lost in the vertebrate ancestor. Similar to *HPGs*, amphioxus *Hox* genes exhibit a temporal colinearity of expression pattern, with the anterior genes expressed in earlier developmental stages than the posterior genes (**Supplementary Fig. S24**).

One major molecular mechanism that contributed to the gene acquisition of amphioxus is segmental duplications, which tend to be of more recent origin and often species-specific (**Supplementary Fig. S25**). Segmental duplications accounted for a higher percentage of the genome in amphioxus vs. vertebrates (9% vs. 3.5%, **Fig. 3f**); they are on average 7.8 kb long, but can be up to 300 kb (**Fig. 3g, Supplementary Fig. S26**). These duplicated segments encompass genes that are enriched for GO categories of G-protein coupled receptor activities, protein tyrosine kinase activities or nucleic acid binding functions (**Supplementary Table S5**). These genes are also frequently enriched for multi-copy ohnologs in vertebrates (*36–38*). Transcriptional factors or genes involved in early development that are often retained after vertebrate WGDs (*39, 40*), however, are not enriched in amphioxus segmental duplicates.

### Developmental dynamics of amphioxus chromatin architecture

Eukaryotic genomes are folded into (active/A or inactive/B) chromatin compartments and to a finer scale of TADs. Such hierarchical three-dimensional (3D) chromatin architectures were previously shown in *Drosophila*, teleosts and mammals to be gradually established or reprogrammed during embryonic development (*41–43*).

To examine whether this is a broadly conserved feature between invertebrates and vertebrates, we collected time-series population Hi-C data of Bf spanning six developmental stages of 1-cell zygote, 32-cell, 64-cell embryos, gastrula, larvae, and adult muscle tissues (**Supplementary Table S9**). Both the percentage of actively transcribed genes (**Fig. 4a**) and the total number of TAD boundaries (TAB) (**Fig. 4b, Supplementary Fig. S27-28**) display a significant (*P* < 0.01, Wilcoxon test) increase after zygotic genome activation (ZGA) around the 64-cell stage (*44*). The strength of TABs measured by insulation scores also becomes generally intensified during development particularly in those strong TABs (**Fig. 4c**). These patterns are similar to those found in Drosophila and mammals (*42, 45*) where major TAD structures of zygote genomes emerge after, although do not necessarily depend on ZGA. In contrast to mammals and Drosophila, the amphioxus genome is highly compartmentalized before ZGA. The A/B compartment strength further becomes significantly (*P*<0.05**, Supplementary Fig. S29**) increased after embryonic stages, but becomes decreased, i.e., possibly reprogrammed on some chromosomes at the gastrula stage (**Fig. 4d-e, Supplementary Fig. S30**).

**Figure 4.**
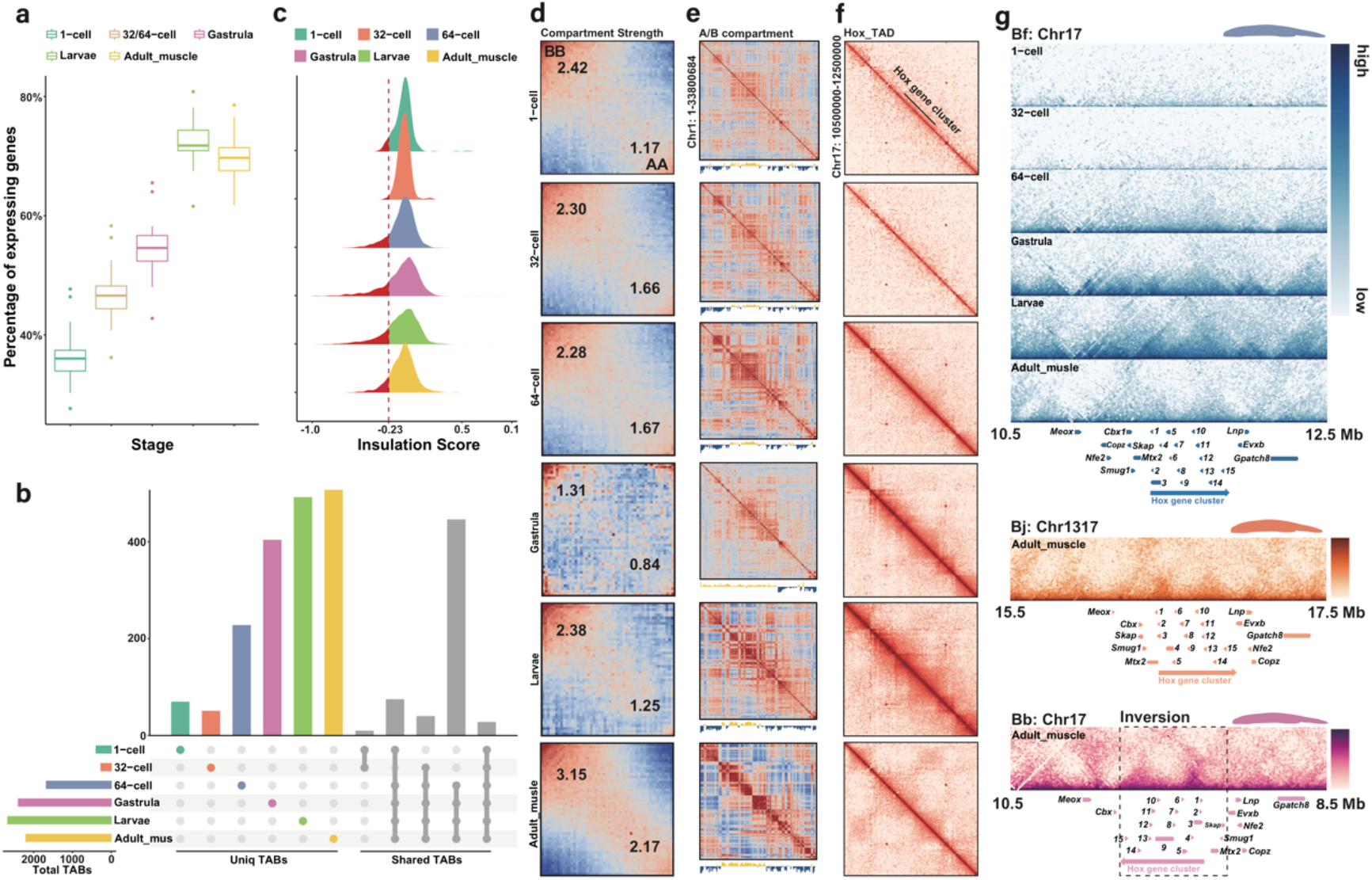
Developmental dynamics of amphioxus chromatin architecture a) The percentage of actively transcribed genes (TPM>1) across five developmental stages of 1- cell zygote, 32/64-cell, gastrula, larvae, and adult muscle tissues of Bf. **b)** The number of TAD boundaries (TABs) at 5kb resolution across six developmental stages of Bf. The horizontal bars show the number of TABs of each stage. The vertical colored bars show the number of specific TABs of each stage, and the grey bars show the number of shared TABs among 6 stages. **c)** The distribution of insulation scores of TABs across different stages. The smaller the insulation score is, the higher strength the TAB has. **d)** Saddle plots of amphioxus Hi-C data binned at 250kb resolution at six different developmental stages. Bins are sorted by their PC1 value. B-B (inactive-inactive) interactions are in the upper left corner, and preferential A-A (active-active) interactions are in the lower right corner. Numbers in the corners show the strength of AA interactions as compared to AB interaction and BB interactions against BA interactions. **e)** Correlation matrix and eigenvector 1(PC1) values value tracks for amphioxus chromosome 1 at 250kb resolution at six different developmental stages. **f)** Distribution of interaction at 15kb resolution at the Bf *Hox* cluster. **g)** Distribution of TADs at the 15kb resolution in three different amphioxus *Hox* regions with the gene tracks.

To explore the formation mechanisms of TADs in amphioxus, we examined the TABs and found that they are enriched for putative binding motifs of chromatin architectural protein CTCF (**Supplementary Fig. S31**), whose transcription level is also specifically increased at ZGA (**Supplementary Fig. S32**). There are disproportionately more (>52%) CTCF-binding site pairs present with convergent forward and reverse orientations at the two TABs of the same TAD (**Supplementary Fig. S33**) (*46*). These results together suggested that similar to vertebrates, loop extrusion facilitated by CTCF protein might play an important role during the formation of TADs upon ZGA of amphioxus. Another mechanism of TAD formation, i.e., self-organization likely mediated by heterochromatin interactions, could also play a role, however it requires chromatin profiling data of different embryonic stages before and after ZGA to be tested in future.

Once established at 64-cell stage, 26.82% of the TABs are overlapped with those in all the later developmental stages, with about 16.88% to 22.95% of TABs only present in one certain stage or tissue (**Fig. 4b**). This indicates that similar to Drosophila and mammals, substantial numbers of, but not all TADs become stabilized and conserved across stages after ZGA, with many others showing dynamic changes during development. To further illustrate this process, we scrutinized the *Hox* cluster of Bf, which is encompassed in one single TAD from 1- to 64-cell stages but becomes segregated into two TADs (**Supplementary Fig. S28**) since the gastrula stage during later development (**Fig. 4f**). The TAB within the *Hox* cluster is weak at gastrula and larvae stages, but becomes clearer in the adult tissue (**Fig. 4f**). This is in contrast to the previous result that characterized the *Hox* cluster of European amphioxus (*B. lanceolatum*) as one TAD, with pooled samples of different embryonic stages and 4C technique (*8*). The *Hox* TABs in adult muscles seem to be conserved across different amphioxus species around *Hox7*. Interestingly, the entire *Hox* cluster of Bb (together with three neighbouring genes) is included in a large genomic inversion (**Fig. 4g**) that occurred after its divergence from Bj in the last 50 MY, with its functional impact on the Bb genome remained to be elucidated in future.

### Evolutionary turnovers of sex determining regions between amphioxus species

The sex-determination (SD) mechanisms of amphioxus remain largely enigmatic, with no cytogenetic evidence for the existence of differentiated sex chromosome pair in Bf and Bb (*19, 47*). A recent genetic study suggested that Bf has a female heterogametic sex chromosome system (male ZZ, female ZW) (*48*). We confirmed this by generating a heterozygous female mutant strain of *Pitx* with transcription activator-like effector nucleases (TALENs), whose mutant alleles are only carried by their daughters. While the mutant alleles can be found in both sons and daughters of male heterozygous mutant strain (**Supplementary Fig. S34**). Using whole-genome re-sequencing data of between 10 to 48 individuals per sex per species (**Supplementary Table S6**), we identified the sexually differentiated regions (SDR) that harbor female-associated SNPs, i.e. excessive female heterozygotes, and are not shared between the three amphioxus species (**Fig. 5a-c**). In particular, the SDR of Bf is located on Chr16 and harbors 189 genes (**Supplementary Table S7**); and those of Bj and Bb are located at two different genomic loci of Chr3, harboring 35 genes and one gene respectively (**Supplementary Table S8**). These SDRs consistently exhibit the highest levels of population differentiation (measured by F_st_) between sexes throughout the genome (**Supplementary Fig. S35**), but do not exhibit sexually differentiated patterns of mapped read coverage. These results together indicated that all three amphioxus species have non-homologous female heterogametic sex chromosomes that have not become differentiated in their genomic sequences.

**Figure 5.**
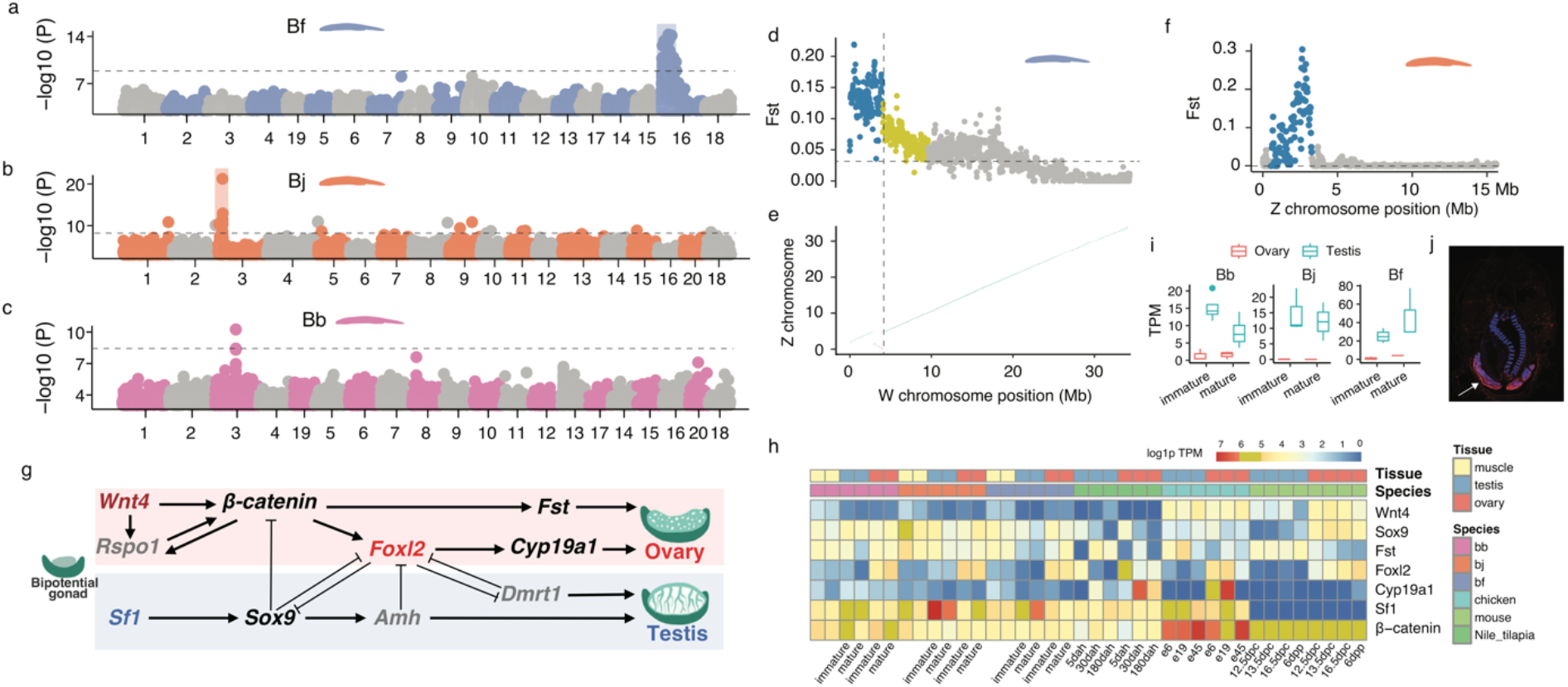
Turnovers of sexually differentiated regions between amphioxus species a-c) Genome-wide association study (GWAS) identified the sex-linked regions in amphioxus. The Y axis shows the log10 transformed p-value of GWAS. **d)** The F_ST_ statistics between male and female populations of Bf reveal the evolutionary strata. Each dot represents a 50 kb sliding window. The horizontal dashed lines show the genomic average levels. **e)** The synteny plot between the Z and W chromosomes of Bf. The purple lines represent reversed alignments. The vertical dashed line indicates the boundary of stratum 1 as well as the inversion. **f)** The F_ST_ statistics between male and female populations of Bj. **g)** The 10 conserved vertebrate SD pathway genes, genes in grey are absent in amphioxus. Only *Foxl2* and *Sf1* are sex-biased in amphioxus. **h)** The expression profiles of chordate SD-related genes over developing gonads. **i)** The candidate Bb SD gene has a conserved testis-biased expression. **j)** RNA fluorescence *in situ* hybridization shows the candidate Bb SD gene has a specific expression in testis

The homomorphic sex chromosomes of amphioxus are similar to those of many fish and frog species, sharing the feature of rapid evolutionary turnovers between species (*49*). This is in contrast to the relatively stable and highly differentiated sex chromosomes of most birds and mammals and may be explained by the ‘fountain-of-youth’ hypothesis. It postulates that occasional sex reversal may induce rare recombination between sex chromosomes and prevent them from becoming differentiated (*50*). Supporting this, we found between 10 to 40% of the phenotypic female or male individuals of the three species exhibit a genotype of the opposite sex in their SDRs (**Supplementary Fig. S36-38**).

With the advantage of fully assembled sequences of ChrZ of Bb and Bj, and particularly those of both ChrZ and ChrW of Bf (**Fig. 1**), we further reconstructed the evolutionary history of these species’ SDRs. The SDR of Bf can be divided into two regions which likely have suppressed or reduced homologous recombination between ChrZ/W at different time points (termed ‘evolutionary strata’ (*51*)). The older stratum spans 4.1 Mb sequence at one end of Bf ChrW chromosome and exhibits uniformly much higher levels of ChrZ/W pairwise sequence divergence and intersexual F_st_ than the rest SDR (**Fig. 5d, Supplementary Fig. S36**). The boundary of this stratum aligns with that of chromosomal inversion between ChrZ/W of Bf (**Fig. 5e**), which probably accounted for the recombination suppression in this stratum. In contrast, the F_st_ values and ChrZ/W sequence divergence levels are not uniform in the rest SDR of Bf (4.1Mb- 11.5Mb), suggesting homologous recombination may have been gradually reduced without involving chromosomal inversions (**Supplementary Fig. S36**). The SDRs of Bj and Bb do not exhibit a pattern of ‘evolutionary strata’ and seem to have gradually reduced recombination, suggested by their F_st_ patterns (**Fig. 5f, Supplementary Fig. S37-38**).

The SDR of each amphioxus species is expected to harbor respective upstream sex-determining genes, which may constitute the sex-determining pathways together with genes on the other chromosomes. We examined the orthologs of 10 reported vertebrate sex-determining genes, and found none of them are present in SDRs of amphioxus. Three upstream SD genes of some vertebrates, *Dmrt1* (**Supplementary Fig. S39**), *Amh* and *Rspo1* do not have an ortholog in the amphioxus genome (**Fig. 5g**); among the rest, only *Sf1* and *Foxl2* exhibit a testis- or ovary-biased expression pattern in amphioxus (**Fig. 5h**). Among the amphioxus SDR genes, we identified a candidate Bj SD gene that is absent in Bf and Bb (**Supplementary Fig. S40**), and a candidate Bb SD gene that are present in Bf and Bj, both of which have specific or biased expression in the gonads, and do not have a vertebrate homolog (**Fig. 5i-j**). These results together indicated that amphioxus and vertebrates independently evolved their SD pathways.

### Conclusions

With three reference-quality genomes of amphioxus, we uncovered their interspecific diversities of genes and chromosomes to an unprecedented resolution. This enabled more direct and accurate reconstruction of ancestral status of the ancestors of both amphioxus and chordates, which was previously based on the draft genome of one amphioxus species. We inferred that there were 20 ancestral linkage groups in the ancestor of *Branchiostoma* amphioxus, best approximated by the Bb genome; and confirmed there were 17 ancestral linkage groups in the chordate ancestor (*13, 16*). Phylogenetic analyses of vertebrate ohnologs and their amphioxus orthologs dated the timing of WGDs, and further characterised the rearrangements and asymmetric loss/retention among the duplicated descendants of CLGs that gave rise to the vertebrate ancestral karyotype. These evolutionarily distant comparisons between amphioxus and vertebrates can be attributed to the slow-evolving genomes of the former relative to those of urochordates.

Our analyses also revealed shared or independently evolved genomic features of amphioxus and vertebrates. For example, both clades seem to establish their major TAD architecture after ZGA, and form two TADs within the *Hox* gene cluster, suggesting these patterns probably originated in their chordate ancestor. In the absence of WGDs, amphioxus species expanded their gene repertoire by segmental duplications or individual gene duplications; and independently evolved their sex-determination pathways from each other, and from vertebrates. By the development of rich genomic resources from this and previous works (*12, 16, 17*), as well as that of gene knockout techniques (*52*), we expect the resurgence of interest into this classic evo-devo model organism, with more functional insights into its genes to be uncovered in future.

## Methods

### Genome sequencing and assembly

Bb and Bj were collected from Xiamen Rare Marine Creature Conservation Areas (Fujian, China) and Bf was introduced from Dr. Jr-Kai Yu’s laboratory (Institute of Cellular and Organismic Biology, Academia Sinica, Taiwan) (**Supplementary Fig. S1**). All of them were cultured as previously described (*52, 53*) . Interspecific hybrids were produced by pooling the sperm of one species, and the eggs of another species except that Bj and Bb cannot be crossed with each other. We extracted high molecular weight genomic DNAs from the muscle tissues of a single individual (male Bj/Bf F1 offspring, Bb/Bf F1 offspring with unidentified sex) using the DNeasy Blood & Tissue Kit (QIAGEN, Valencia, CA), and inspected the DNA quality by Qubit 2.0 Fluorometer (Thermo Fisher Scientific, Waltham, MA) and 2100 Agilent Bioanalyzer (Agilent). We prepared the 20 kb SMRTbell^TM^ PacBio libraries and generated sequencing data of ∼50G for the two hybrids (Bb/Bf and Bj/Bf). We estimated the heterozygosity levels of three species’ genomes using Illumina reads of the three species using GenomeScope (*54*). For the hybrids, the estimated genome size was equivalent to the sum of the haploid genome sizes of the parental species (**Supplementary Fig. S1**).

We used Falcon (*55*) to assemble the PacBio subreads of two hybrid samples, after discarding raw subreads and corrected reads (preads) shorter than 8kb. We used the following parameters to avoid collapse of reads derived from different parental species: pa_HPCdaligner_option = -v -dal128 -t8 -e0.75 -M24 -l3200 -k18 -h480 -w8 -s100, ovlp_HPCdaligner_option = -v -dal128 -M24 -k24 -h1024 -e.96 -l2500 -s100. We used the arrow (from the Falcon assembler) algorithm to polish the contigs twice, followed by another two-round polishing with the Illumina reads derived from the same hybrid individual, using pilon (1.22) (*56*). To assign the contig sequences of hybrids to each parental species, we aligned the Illumina reads of either parental species to the contigs by bwa-mem with default parameters, and only kept the alignments with a mapping quality higher than 60. For each contig, we calculated the proportion of nucleotide sequences that were mapped by each species’ reads (coverage), without considering the contigs shorter than 20kb. We assigned a contig to either parental species if the sequencing coverage was larger than 10% for one parental species, while the sequencing coverage for the other species was below 1% (**Supplementary Fig. S2**). We then used minimap2 (2.15-r905) (*57*) to align the PacBio reads of hybrids to the assembly, with the option ‘--secondary=no’, and partitioned the species-specific haploid reads. These partitioned reads were used for assembling the four haploid assemblies (one Bb, one Bj and two Bf) by Canu (1.6) (*58*) (‘corOutCoverage=200 correctedErrorRate=0.15’) and Falcon (‘pa_daligner_option= - k18 -e0.7 -l2000 -h480 -w8 -s100, ovlp_daligner_option=-k24 -e.93 -l2000 -h600 -s100’). Since the read length of Bb/Bf was longer, we increased the ‘-l’ parameter from 2000 to 2500 in ‘pa_daligner_option’ and from 2000 to 3000 in ‘ovlp_daligner_option’. The polishing steps were similar to those for the diploid assembly of hybrids. Then contigs of two pipelines were merged: we aligned the Canu contigs against the falcon contigs using the nucmer aligner (MUMmer 3.0) (*59*) with the option -b 400. When one Falcon contig spanned the boundaries of two Canu contigs, we linked the Canu contigs with a gap of 200 Ns.

Finally, we used the Juicer (1.7.6) (*60*) pipeline and 3D-DNA (180922) (*61*) to connect the contigs into chromosome-level scaffolds. To reduce the false-positives of contig splitting, we used the following parameters: --editor-coarse-resolution 500000 --editor-coarse-region 1000000 --editor-saturation-centile 1 -r 0 --editor-repeat-coverage 1 --editor-coarse-stringency 70. We manually curated the chromosome assembly by editing the Hi-C contact map using Juicebox (1.90) (*62*). After that, we updated the assembly using the ‘review’ module of 3D- DNA. The unanchored scaffolds are highly repetitive, with repeat content as high as 79.0%, 63.5% and 81.7% for Bb,Bj and Bf respectively.

### Genome annotation

To annotate genes, we generated Iso-seq and RNA-seq data from whole-body adult male and female individuals of the three species. We used IsoSeq3 (3.1.0) (*63*) and Trimmomatic (0.36) (*64*) for pre-processing the raw reads. Then we generated reference-guided and *de novo* assembled transcript sequences using Cupcake (5.8) with Iso-Seq reads, and StringTie (1.3.3b)(*65*) (-m 300 -j 5 -c 8) and Cufflinks (2.2.1) (*66*) (–multi-read-correct –max-intron-length 30000) and Trinity (2.6.6) (*67*) (--min_glue 10 --path_reinforcement_distance 30 -- min_contig_length 400 --jaccard_clip) with RNA-seq reads. We then used the Mikado (1.2.2) (*68*) to integrate all transcript sequences. We used RepeatModeler (1.0.10) (*69*), Tandem Repeat Finder (409) (*70*) (‘2 7 7 80 10 50 500 -d -l 6’) and MITE_Hunter (*71*) (‘-I 86 -n 8 -c 8’) for annotating and classifying the repeat families.

To produce a consensus gene model, we ran MAKER (2.31.10) (*72*), after masking the annotated repeats. We used the query protein sequences from NCBI RefSeq database (Bb: GCA_001625405.1 and Bf: GCA_000003815.1), and the transcriptome annotations produced by Mikado. The MAKER gene annotation was then used to train SNAP (2013-11-29) (*73*) (maker2zff -c 0.99 -e 0.99 -o 0.99 -l 800 -x 0.01) and AUGUSTUS (3.3) (*74*) for *ab initio* predictions. Gene evidence from protein alignment, StringTie transcripts, ISO-seq transcripts, SNAP and AUGUSTUS predictions, were combined by EVidenceModeler (EVM) (1.1.1) (*75*), with the highest weight on the protein alignment and StringTie transcripts (10), intermediate weight on ISO-seq transcripts (5) and lowest weight on the *ab initio* predictions. We used the PASApipeline (v2.3.3) (*76*) to polish the gene models. We used InterProScan (5.35-74.0) (*77*) to annotate gene ontology (GO) for the predicted coding genes.

To annotate putative centromeres, we counted the copy number and total length for each satellite repeat based on the RepeatMasker results and inferred the most abundant and longest satellite sequences to be associated with centromeres. The identified centromeric monomer of Bf is consistent with the reported result (*78*). The recombination rates were estimated with ReLERNN program, using the individually sequenced data (**Supplementary Table S8**). The nucleotide diversity was estimated in 100 kb windows using VCFtools (0.1.16) (*79 80*). To verify the centromere with the fluorescence in situ hybridization (FISH) technique, we prepared the probe of candidate centromeric monomer and the slides of Bf chromosomes. The details of the FISH experiment were described in Lie et al. (2002) (*79*). To annotate telomeres, we searched for clusters of (AACCCT)n repeats throughout the genomes using RepeatMasker. We only kept those with a total length of 200 bp (33.3 consecutive AACCCT repeats) to reduce false positives. We used the R package Quadron (*81*) to predict the G-quadruplexes (G4) throughout the genome with default settings. Then we calculated the length of G4 elements over 20kb sliding windows along the chromosomes using bedtools coverage.

### Comparative genomic analyses

We included three amphioxus species and four vertebrate species to infer the orthologous gene groups. The Refseq annotations of human (GCF_000001405.39), mouse (GCF_000001635.26), zebrafish (GCF_000002035.6) and chicken (GCF_000002315.6) were downloaded from NCBI. When multiple isoforms were present, we selected the longest one for each gene. We ran OrthoFinder (2.2.7) (*82*) to group the orthologous genes, using diamond (0.9.21) for protein alignment. We used Last (1042) (*83*) to align genomes of mouse (GRCm38.p4), chicken (GRCg6a), zebrafish (GRCz11), Bb, Bj and Bf against the human reference genome (GRCh38.p12), with -uMAM4 for mouse alignment, and more sensitive -uMAM8 for the other species. The one-to-one best alignments were retained and merged by Multiz (v11.2) (*84*).

For the reconstructing the chordate phylogeny, we added *Ciona intestinalis* (GCA_009617815.1) (*85*) and scallop (*Mizuhopecten yessoensis*, ASM211388v2) (*86*), with the latter set as an outgroup. We excluded the alignments in which the sequences were aligned to non-homologous chromosomes among amphioxus, because they likely represent alignment errors. The filtered alignments contained 5,074 loci, with a total size of 276,373 bp. We used IQ-TREE (2.0-rc1) (*87*), with the substitution model (TVMe+R3), to construct the phylogenomic tree, and ran bootstrapping for 100 times.

We used the PhastCons from the PHAST package (1.5) (*88*) to annotate the conserved non-coding elements across the genomes. First we used (msa_view --4d) all fourfold degenerate (4d) sites for estimating a nonconserved phylogenetic model by phyloFit (a PHAST program), with the phylogenetic tree as ((((human,mouse),chicken),zebrafish),(bf,(bb,bj))). Then we ran the PhastCons program with the alignments and the nonconserved model to estimate the rho value and the conserved model (the nonconserved model remained the same as the 4d model). Finally, we ran the PhastCons program again but added --most-conserved option to identify the conserved elements. Then we compared the conserved elements with annotated features of the human genome (RefSeq GCF_000001405.39 and Ensembl annotation) using BEDTools (2.29.0) (*89*). For each conserved element, we assigned it to one feature if overlapped. If multiple features were overlapped with a single element, they were assigned under the following priority: protein-coding region > pseudogene > non-coding RNAs > lncRNA > UTR > intron > intergenic.

### Ancestral karyotype reconstruction

We generated whole genome alignments between amphioxus species by minimap2 (2.15-r905) (*58*) (-x asm20) and visualized the alignments by D-Genies online tool (*90*) (**Supplementary Fig. S14**). We selected 7269 orthologous gene groups (orthogroups) in which Bb genes are located within the same chromosome. 1799 orthogroups contained more than one gene in chicken which were informative for reconstructing the chordate ancestral karyotype. For each Bb chromosome (*i*), we asked which chicken chromosome (*j*) its homologous genes belong to, and counted the gene number for each chicken chromosome (CK*ij*). Then we calculated the relative abundance of genes of a chicken chromosome for a given Bb chromosome (nCK*ij*):

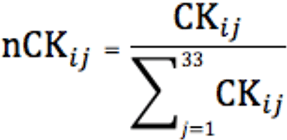

We included 33 chicken chromosomes, and retained a chicken chromosome when the nCK*ij* value was larger than 4%, for a given Bb chromosome (*i*). Then we visualised the nCK*ij* values for every Bb chromosome with a network-style graph (**Fig. 2b, Supplementary fig. 16**), using the igraph R package. We used 244 orthogroups that retained three or four chicken ohnologs and performed coding sequence alignments using MAFFT (v7.294b) (*91*). Then we constructed the phylogenetic tree using concatenated sequence alignments of the same CLG using IQ-TREE, with 1000 times bootstrapping. Based on the phylogenetic relationships, we assigned the four ohno-chromosomes derived from a single CLG as ohno-A, ohno-B, ohno-C and ohno-D. For each ohno-chromosome group, we further included the orthologous genes of human, mouse and spotted gar of the chicken gene in that group (**Supplementary Table S3** and **Supplementary Fig. S20**) Then all the coding sequences of three amphioxus species and vertebrates were aligned with MAFFT (7.427) (*91*) and GUIDANCE2 (2.02) (*92*) pipeline, producing concatenated alignments with 409,659 nucleotide sites. We then used BASEML (4.9j) (*93*) to estimate the overall mutation rate with the time calibration on the root node (550 MY for the vertebrate and amphioxus split (*94*)). The topology “((bf,(bb,bj)),((((chicken-Ohn_A,(human-Ohn_A,mouse- Ohn_A)),gar-Ohn_A),((chicken-Ohn_B,(human-Ohn_B,mouse-Ohn_B)),gar- Ohn_B)),(((chicken-Ohn_C,(human-Ohn_C,mouse-Ohn_C)),gar-Ohn_C),((chicken-Ohn_D,(human-Ohn_D,mouse-Ohn_D)),gar-Ohn_D))));” was used. General reversible substitution model and discrete gamma rates were estimated by maximum likelihood approach under the strict clock. The divergence time was then estimated using MCMCtree (4.9j) (*95*), with three soft-bound calibration time points: 534–566 MY for the vertebrate and Branchiostoma species split, and 62-101 MY for the human and mouse split, and, 306-332 MY for the chicken and mammal split, 416-422 for the teleost and tetropad split (*96*).

### Gene evolution

We used SDquest (0.1) (*97*) to identify segmental duplications (SDs) in all amphioxus species and four vertebrates including human (hg38), mouse (mm10), chicken (galGal6) and zebrafish (danRer11). We excluded the sex chromosomes of human, mouse and chicken, and alternate-loci scaffolds of zebrafish as these sequences may confound the identification of SD. We only kept SDs that are longer than 1000 kb, and show a sequence similarity level of at least 70%. For studying gene gain and loss, we selected 8,464 orthologous gene groups that contain at least one vertebrate species and one amphioxus species as the input for Notung (2.9.1) (*98*) gene family reconstruction. We identified 200 orthogroups that had more than one gene copy in all amphioxus species, but had single-copy genes in vertebrates. The mean copy number of the expanded gene families were 3.6, 3.8 and 4.8 for Bb, Bj and Bf respectively.To elucidate the evolution of the *Hox* genes across chordate species, protein and CDS sequences of chicken, mouse, human and zebrafish *Hox* genes were downloaded from NCBI, and aligned to those of amphioxus species by MAFFT (v7.407), with alignment polishing by trimAl (v1.4.rev15) (*99*). We used IQ-TREE to infer the phylogeny, and the AVX+FMA model was selected automatically by IQ-TREE. We used EvolView online tool (https://www.evolgenius.info/evolview) to visualize our phylogenetic tree. RNA-seq data of multiple Bf developmental stages were downloaded from NCBI SRA (PRJDB3785) for estimating the *Hox* gene expression level using HISAT2 (2.0.4) and featureCounts (v1.5.2).

### 3D genome analyses

*in situ* Hi-C libraries were constructed from the muscle and embryonic tissues of amphioxus as described before (*100*). Hi-C data were mapped to the genomes using bwa-mem (0.7.17-r1188) with parameters ’-A 1 -B 4 -E 50 -L 0’. The quality control including valid pairs and cis/trans ratio of Hi-C data was finished by using pairtools(0.3.0) ( https://pairtools.readthedocs.io/en/latest/) and the estimated resolution was calculated by HiCRes(2.0) (*101*) Then the mapped read-pairs were used to generate raw Hi-C contact matrix at 5kb, 15kb and 30kb resolution using hicBuildMatrix of the HiCExplorer (2.2.1) suite (*102*). We used the ICE method implemented in hicCorrectMatrix to remove the bins with extremely low or high numbers of reads, and visualized the matrix with hicPlotMatrix. We used hicFindTADs to generate the coordinates of TADs and the TAD insulation score of each bin (-- thresholdComparisons=0.01, --delta=0.01). To investigate the overlaps of TAD boundaries between different developmental stages, we combined the TAD boundaries of all development stages into one set, and extended each boundary for 5kb of both sides to form 15kb windows and merged adjacent windows when their distance was not longer than 10kb. This generated a set of boundaries that existed in at least one developmental stage. We then compared the boundaries of each stage to this common set, and defined conservation of boundaries as an overlap of at least 15 kb in size. We used cooltools (0.3.2)(https://cooltools.readthedocs.io/en/latest/) and its call-compartments function to obtain the first eigenvector values (PC1) of each chromosome of the Hi-C matrices with a 250kb resolution. Regions with positive PC1 values are assigned as A (active) compartments and those with negative PC1 values are assigned as B (inactive) compartments, adjusted by the gene density of the region. Compartment strength was calculated as AA × BB/AB^2^ for each chromosome. Saddle plot was also obtained by cooltools. In brief, enrichment contact maps at the 250kb resolution were normalized by genomic distance into a 50 by 50 bin matrix to calculate the observed/expected (O/E) values as contact enrichment. Bins in the matrix were sorted by PC1 values and all contacts with similar PC1 values were aggregated to obtain compartmentalization saddle plots with B-B interactions in the upper left corner and A-A interactions in the lower right corner. The numbers in saddle plots indicate the strength of the top 20% of A-A interactions (over A-B interaction) and the bottom 20% of B-B interactions (over B-A interactions). We used FIMO (*103*) to search for human CTCT motif (MA0139.1) in the amphioxus genomes and identified 62,987 putative CTCF motifs. To test whether the CTCF motif was enriched in the TAD boundary, we used bedtools intersect to identify the CTCF motifs located in the 15kb TAD boundaries (5kb boundary extended by 5kb of both sides) of the six developmental stages. In addition, we also checked whether the TAD boundaries contain more CTCF motifs than by chance, we randomly selected 15 kb windows across the genome and calculated the proportion of windows that contain CTCF motifs (**Supplementary Fig. S41**). The pairings of convergent CTCF sites at domain boundaries is considered as a hallmark of the conserved role of CTCF/cohesion in TAD formation (*104*). The enrichment pattern of putative CTCF binding sites (**Supplementary Fig. S32**) and the distribution pattern of convergent CTCF site pairs (**Supplementary Fig. S34**) were similar for TAD results derived from different TAD-calling bin sizes.

### Sex chromosome analyses

*Pitx* mutants were generated and detected using the TALEN method as described before (*105*). The TALEN pair used for mutant generation are Fw3: 5’-GCAACCGTTCGACGAC-3’ and Rv3 5’-TGTAGGCCGGCGAGTA-3’ which are from the third coding exon of the gene. A *Tat*Ⅰrestriction site was included in the target site for genotyping and primer pair used for genotyping are Pitx-TALEN-PCR-F2 (5’-AGGTCTGGTTCAAGAACCG-3’) and Pitx-TALEN-PCR-R4 (5’-TCACGGTAAGCGTAAGGCTG-3’). Two different mutant stains were generated. The founder of stain 1 is a female, which was crossed with a wild type male to generate F1 offspring, from which a female heterozygote was further crossed with a wild type to generate F2 descendants. In contrast, the founder of strain 2 was a male and an F1 heterozygous male was used to generate its F2 descendants.

We generated re-sequencing Illumina data of multiple individuals of both male and female (on average 25 individuals of each sex) at a coverage larger than 20X (**Supplementary Table S6**). The raw reads were mapped to the reference genome using bwa-mem (0.7.16a), with default parameters. After sorting the alignments with samtools (1.9) sort, we marked the duplications of reads using the MarkDuplicates function of the picardtools package (2.14.0). We then used the GATK (3.8) (*106*) pipeline to call variants. To do so, we ran HaplotypeCaller to generate GVCF output for each sample. This was done separately for each chromosome with the interval (-l) option, and then the GVCF outputs were combined with the GatherVcfs function of the Picard toolkits (*107*). Then we genotyped the variants with the GVCF files as inputs of all samples (joint calling) using GenotypeGVCFs. We selected single-nucleotide variants (SNPs) for further analysis and filtered the SNPs with the following criteria: QD < 2.0 || FS > 60.0 || MQRankSum < -12.5 || RedPosRankSum < -8.0 || SOR > 3.0 || MQ < 40.0. We used the biallelic SNPs (filtered by bcftools -m2 -M2) to screen for sex-linked variants. We further excluded the variants that have minor allele frequency less than 0.05 and missing rate larger than 10%. We used Beagle (28Sep18.793) (*108*) to do the imputation for the variants and obtain an initial set of phased genotype calls for all variants. SHAPEIT (v2.r904) (*109*) was then used to produce a more accurate set of phased genotypes on the variants. A total of 4,954,852, 7,213,889, and 12,016,687 high-quality phased SNPs in Bj, Bb and Bf respectively, were used to perform whole-genome association analysis for the sex trait (male or female) with EMMAX (efficient mixed-model association expedited, version 8.22) (*110*). Population stratification and the hidden relatedness were modeled with a kinship (K) matrix in the emmax-kin-intel package of EMMAX. The genome-wide significance thresholds of all tested traits were evaluated with the formula P=0.05/n (where n is the effective number of independent SNPs). Apart from identifying sex-associated regions, we screened for differentiated regions between the sexes. We calculated the F_ST_ values between male and female populations using VCFtools (0.1.13) (*80*). SNPs with more than two alleles were removed. The F_ST_ values were estimated in a 10 kb sliding window with an overlapping size of 5 kb. For Bb whose sex-determining region is much smaller, we used 5 kb windows instead of 10 kb. We defined the non-recombining regions of the sex chromosomes by the sex-linked SNPs identified through the whole-genome association tests. We evaluated the extend of sex chromosome differentiation with two measures: 1) F_ST_ and 2) the difference between male and female SNP density.

We collected transcriptomes of immature (identifiable but not functionally mature) and mature gonads for studying the candidate sex-determining genes of amphioxus. We mapped the RNA-seq reads against the genomes using HISAT2 (2.1.0) with the parameters ‘-k 4 --max-intronlen 50000 --min-intronlen 30’. The alignments with mapping quality score lower than 10 were removed (samtools view -q 10). Then we used featureCounts (1.6.2) (*111*) to count the reads mapped to the annotated transcripts. We used the TPM (transcripts per million) method to quantify and normalize the expression levels.

We chose 10 conserved vertebrate SD pathway genes: *Wnt4*, *Sf1*, *β-catenin*, *Rspo1*, *Sox9*, *Amh*, *Foxl2*, *Fst*, *Cyp19a1* and *Dmrt1* to check their presence or absence in the amphioxus genomes. We first checked the orthogroups that contain those SD genes and whether amphioxus is present in these orthogroups. If amphioxus is absent in the orthogroups, we searched the coding sequences of the SD genes against the amphioxus genomes by BLAST **(Supplementary Fig. S40)**. The absence of *Dmrt1* in amphioxus is consistent with a recent study (*112*).

## Author contributions

Q.Z., L.X., Q. J. Z., G. L. conceived the project; Z. H., Y. Z., D. C., S. P., T. X., W. C., C. S., X. W., Y. H., C. X., Y. N.Y., Y. Y., W. H., X. H., Y. Z., Y. C., C. B., C. H., L. X., S. X., Z. Y., Y. J. acquired the data; Z. H., L. X., C. C., J. L., Z. Z., W. K., Q. Z. performed the analyses; Z. H., L. X., C. C., J. K.Y., E.D.J., G. L., G. L., Q. J. Z., Q. Z. wrote the paper.

## Data and code availability

The genome assemblies have been deposited at GenBank under the accession PRJNA603158, PRJNA603159, PRJNA647830. All sequencing data has been deposited at PRJNA602496. A full list of accessions is available in the **Supplementary Table S8**. The scripts used in the study have been deposited at Github (https://github.com/lurebgi/amphioxusGenome)

