## Supplemental figures for "Three amphioxus reference genomes reveal gene and chromosome evolution of chordates"

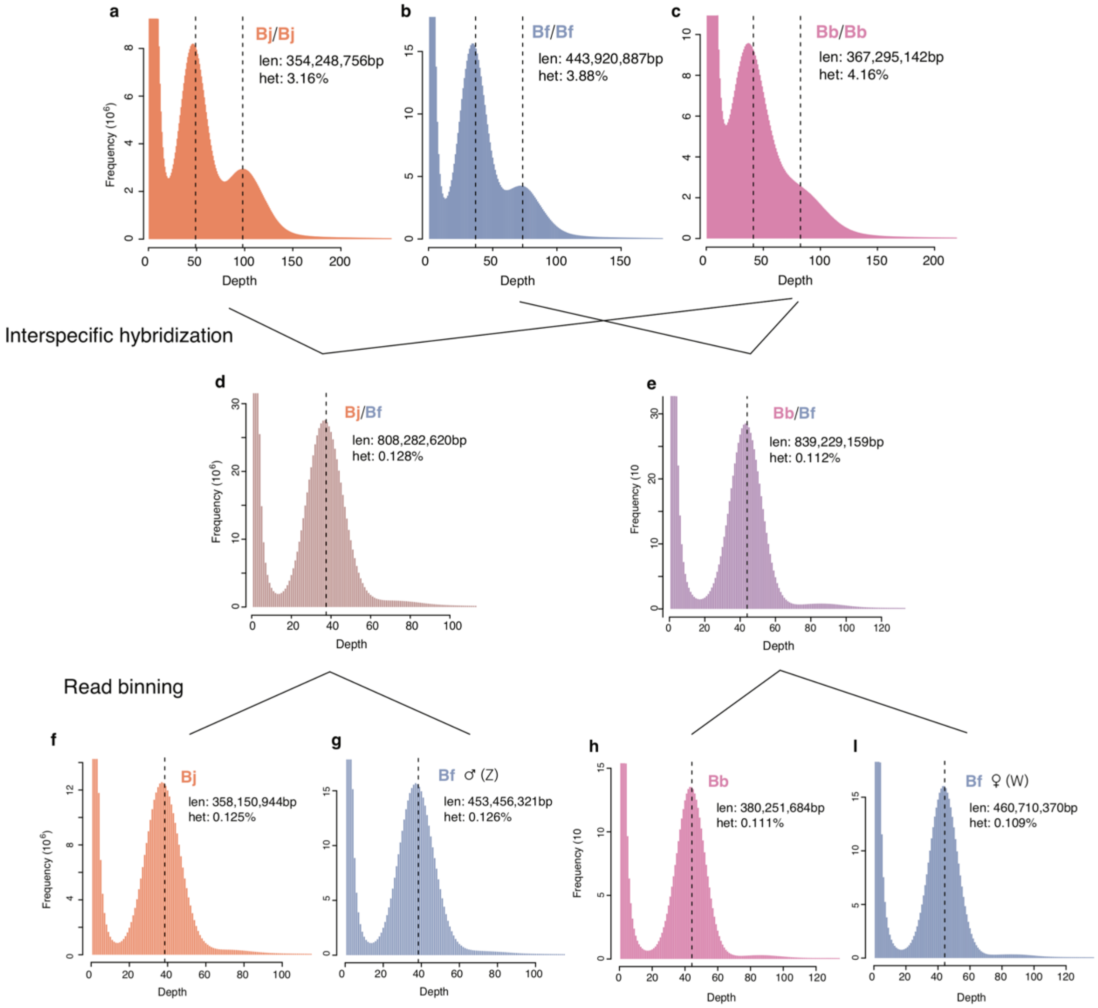


**Supplementary Fig. S1 Estimation of genome size and heterozygosity levels for pure species and the hybrids**.

K-mer size of 21 was used. The distribution of k-mer counts were taken as input for genomeScope to estimate the genome size and heterozygosity of the parental species (**a-c**) and their hybrids (**d-e**). The parental species have one extra peak at the half-coverage level, suggesting the high levels of heterozygosity, while the hybrids have one single peak, suggesting the interspecific hybrid genomes are effectively haploid. After partitioning the species-specific reads from the sequencing data of the hybrids (**f-i**), the species-specific reads show a single haploid peak, the estimated genome size represents the haploid genome size of the pure species.

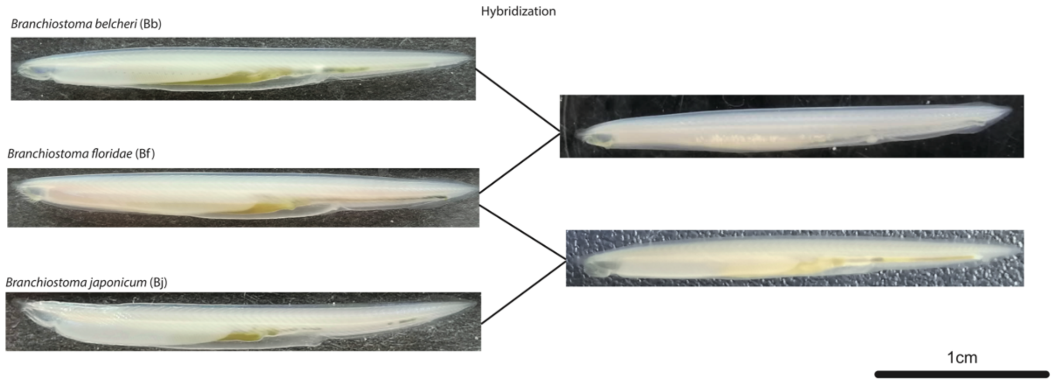


**Supplementary Fig. S2 The pure species of three amphioxus species and their hybrids**.
Photo credit: Zhen Huang, Fujian Normal University

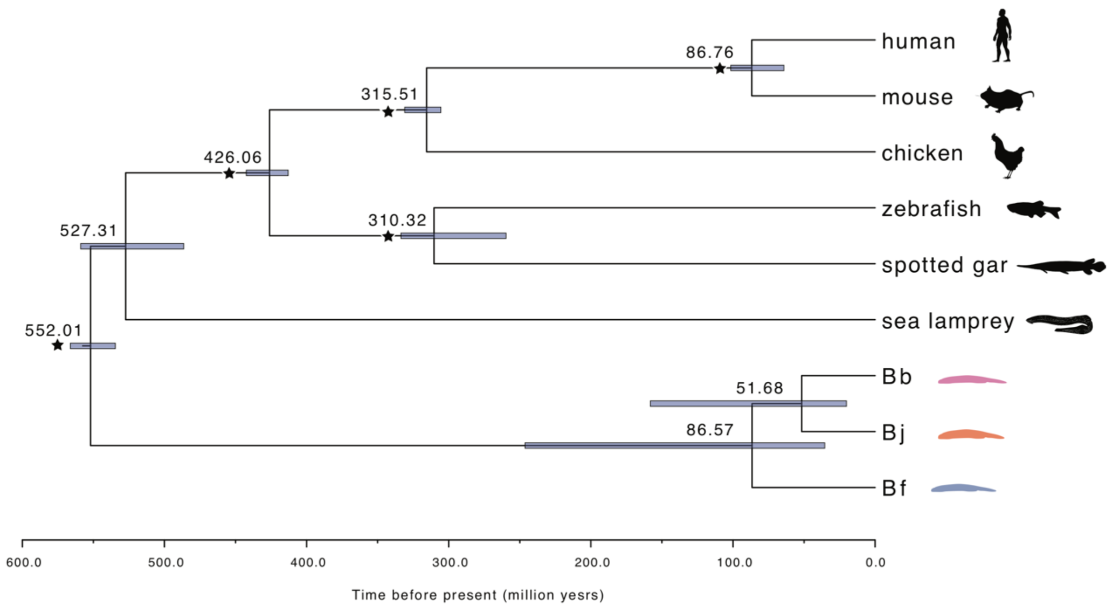


**Supplementary Fig. S3 Molecular dating of amphioxus species**.
The 4-fold degenerate (4D) sites were extracted from the whole genome alignments of four vertebrates and three amphioxus species, according to the annotations of human gene models. The bars show the 95% confidence interval of divergence time. The black asterisks represent the nodes whose divergence time have been calibrated with fossil records.

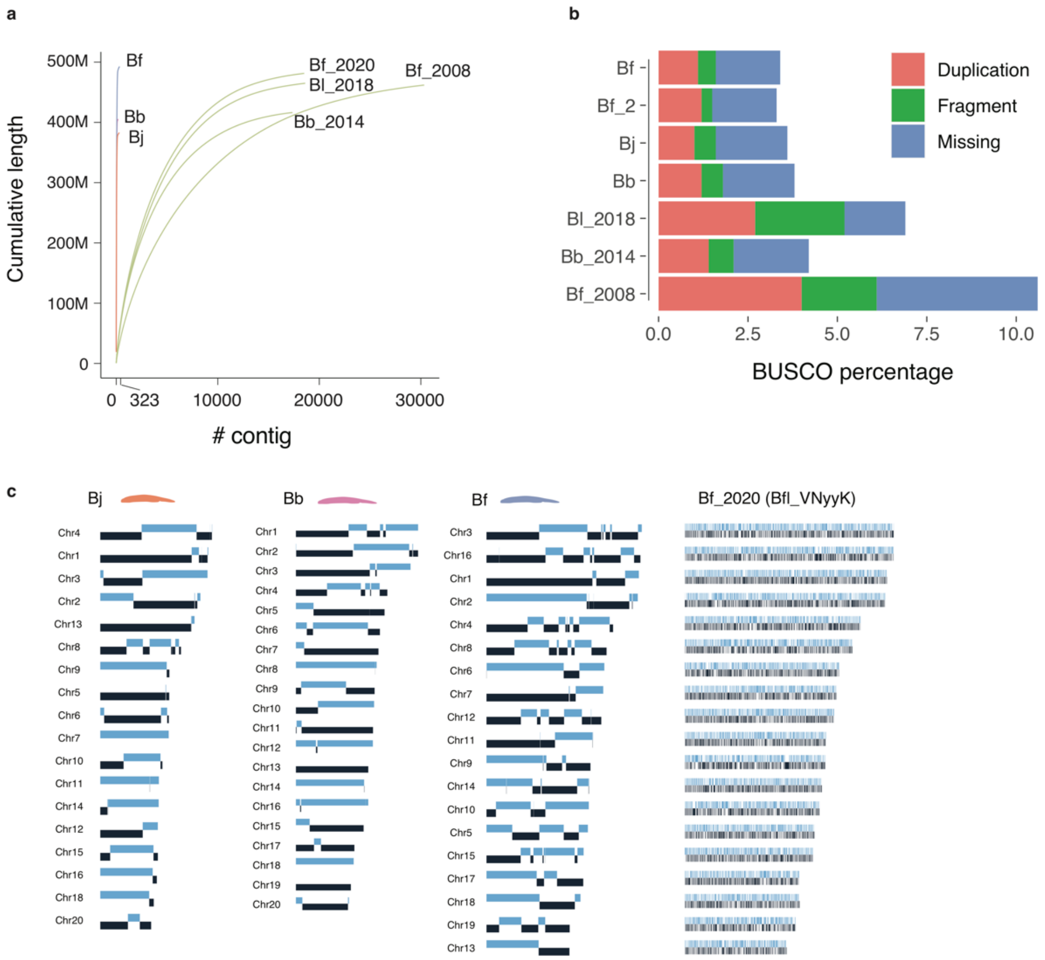


**Supplementary Fig. S4 Improvement of contig length by more than 200 folds**.

**a)** The long-read assemblies produced in this study (Bf, Bb and Bj) have a much longer contig size than the previous short-read assemblies (Bf_2020 GCA_000003815.2, Bf_2008 GCA_000003815.1, Bl_2018 GCA_900088365.1, Bb_2014 GCA_001625305.1). The long-read assemblies have a small contig number (maximum 323), while the short-read assemblies contain at least 16,000 contigs.

**b)** Percentage of BUSCO genes of the duplication, fragment and missing categories. Our new genome assemblies tend to have a lower percentage for those genes.

**c)** The distribution of contigs on the chromosomes. The dark and light blue blocks represent contigs. Bf_2020 (Bfl_VNyyK) refers to the Bf genome published by (Simakov et al. 2020)

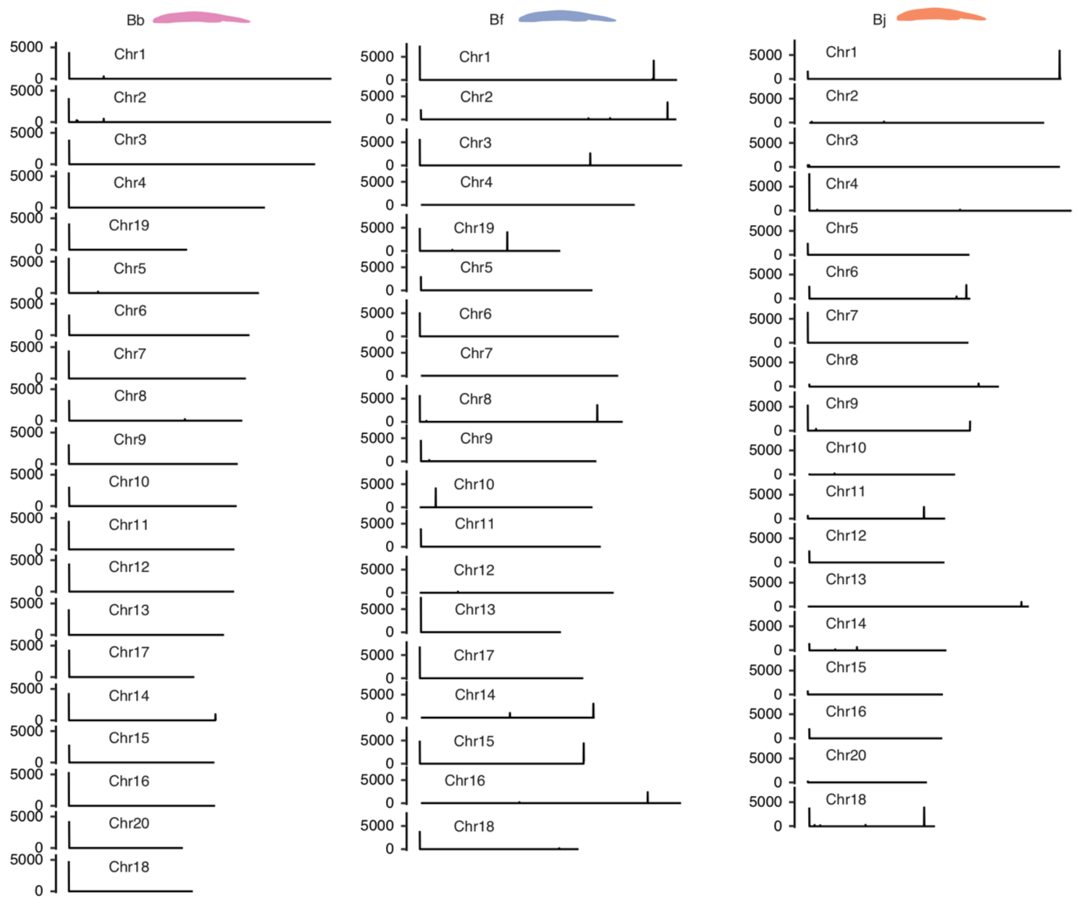


**Supplementary Fig. S5 The distribution of telomere repeats along the chromosomes**. The positions of the telomere repeats (GGGTTA)n were annotated with RepeatMasker. The lengths of the telomere repeats were summed up in every 5 kb window along the chromosomes were plotted. On most chromosomes the telomere repeats are enriched at one chromosome end.

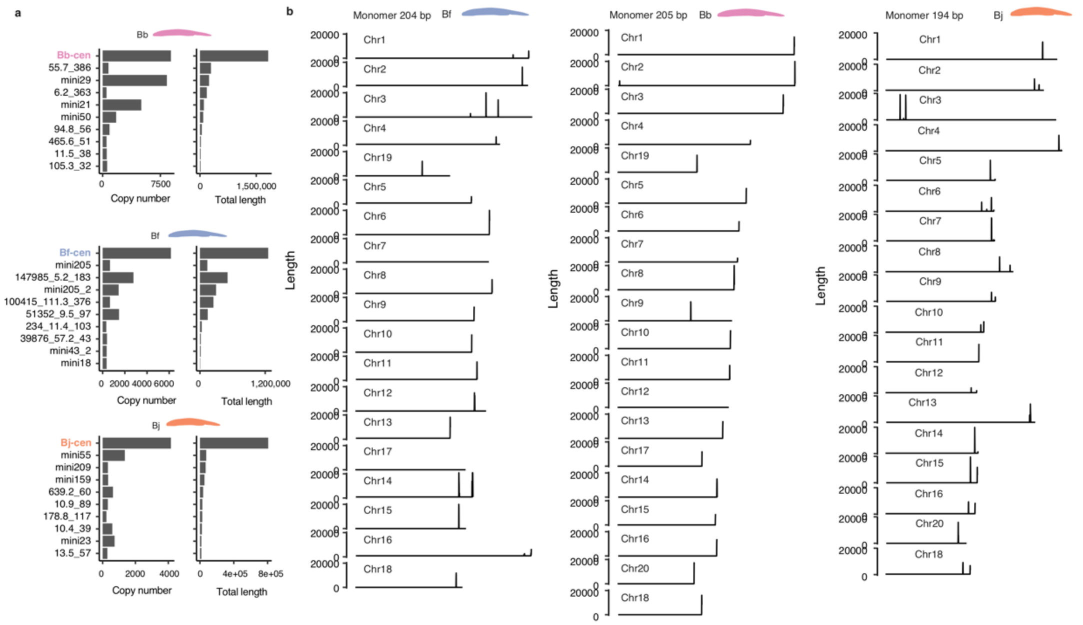


**Supplementary Fig. S6 The identification and distribution of centromeric repeats.**
**a)** The copy number and total lengths of top 10 most abundant satellite repeats. The centromeric satellite repeats (top 1, colored) have the largest copy numbers and longest total lengths.
**b)** The length of centromeric repeats were summed up in every 20 kb window and plotted along the chromosomes. In most cases, the centromere repeats appear as a single locus on the chromosomes.

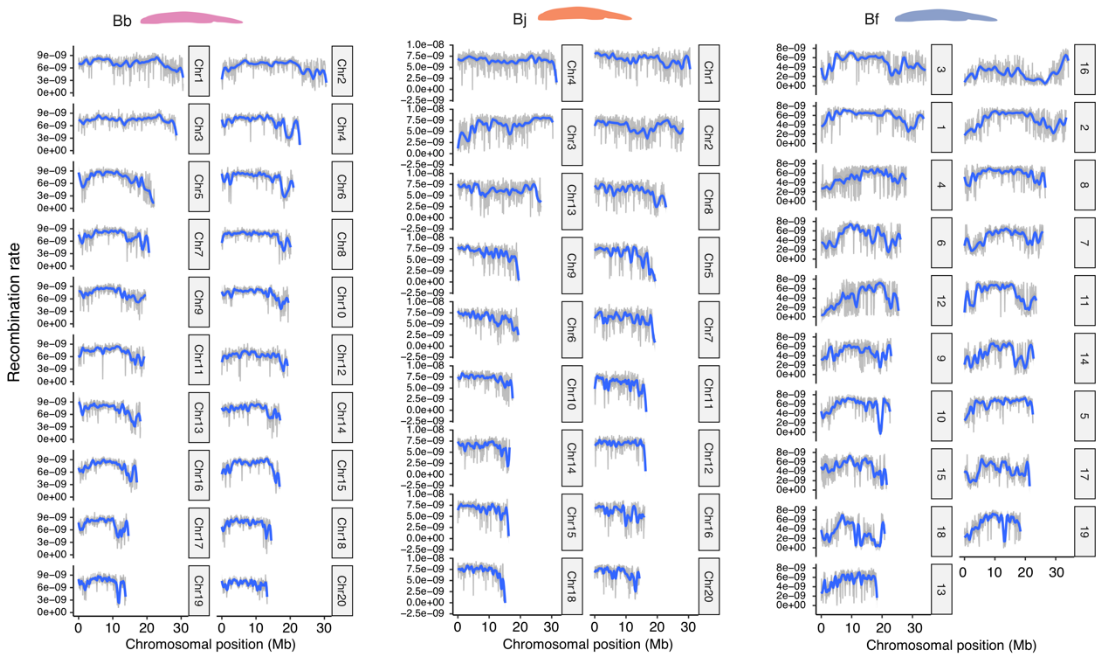


**Supplementary Fig. S7 The low levels of recombination rate at centromeric and pericentromeric regions**.

The recombination rates were estimated with the ReLERNN pipeline, using the genome-wide population data. The grey lines show the original estimates while the blue lines show the smoothed means. In most chromosomes the recombination rates tend to be lower in the regions towards the 3’ ends where the centromeres are located.

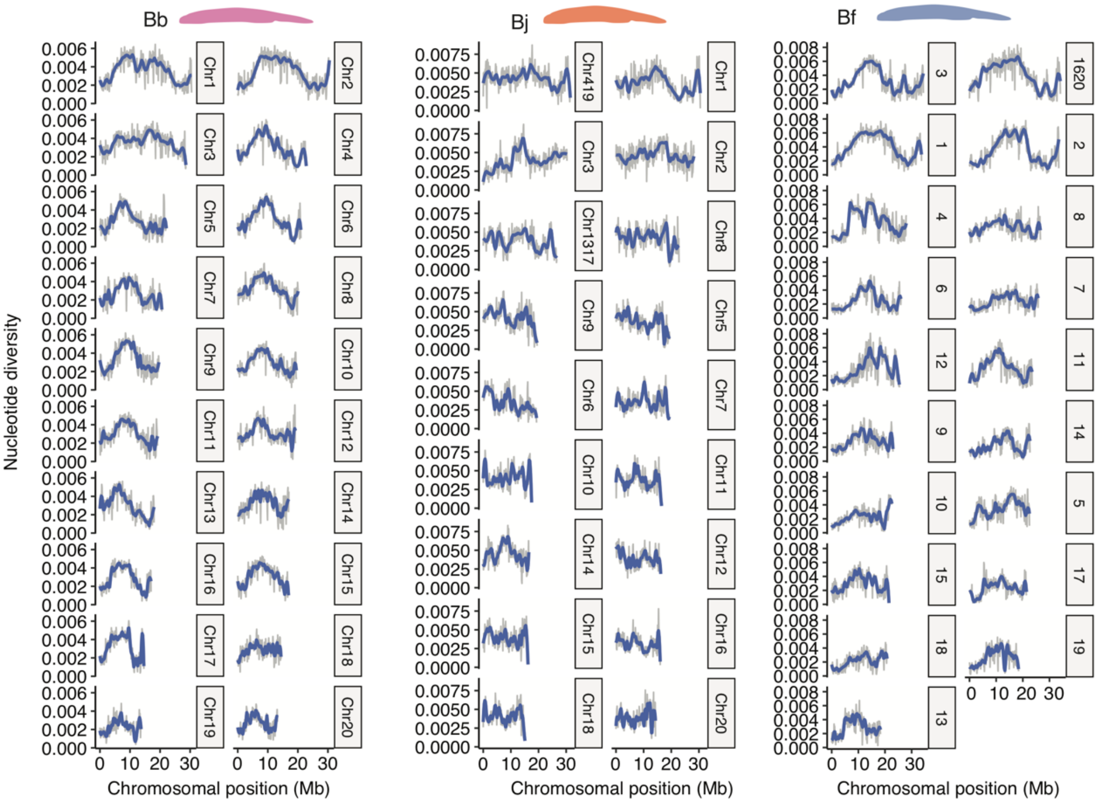


**Supplementary Fig. S8 The low levels of nucleotide diversity at centromeric and pericentromeric regions**.

The nucleotide diversity (π) was estimated using the same population data as in Supplementary Fig. S7. We used VCFtools to estimate the nucleotide diversity in 100 kb windows. The π values tend to drop near the regions close to the 3' ends of the chromosomes.

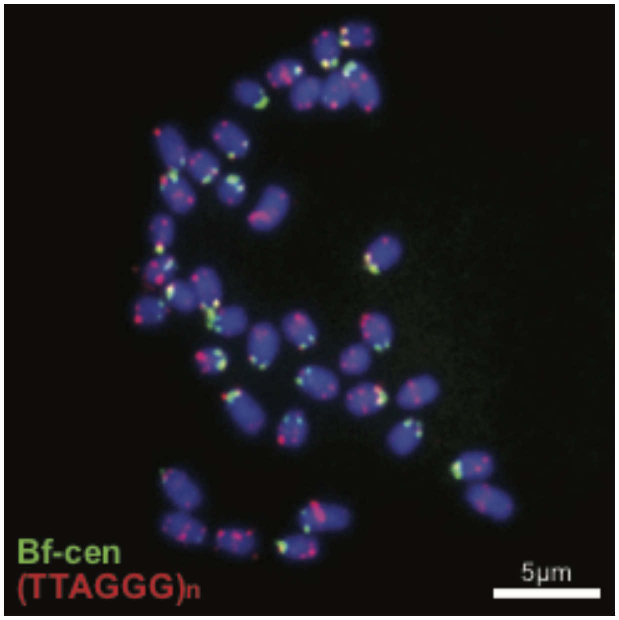


**Supplementary Fig. S9 Fluorescent in situ hybridization (FISH) of centromere and telomere probes to Bf chromosomes**.

The 204-bp Bf centromeric monomer sequence (Bf-cen) is shown in Supplementary Fig. S10. The probe (green) is hybridized mostly to the chromosomal ends. The probe of the telomeric motif (TTAGGG)n (red) is hybridized to the other ends of chromosomes, sometimes colocalizing with the centromeric probes. On some chromosomes interstitial telomeric repeats that appear in the middle of the chromosomes are also detected.

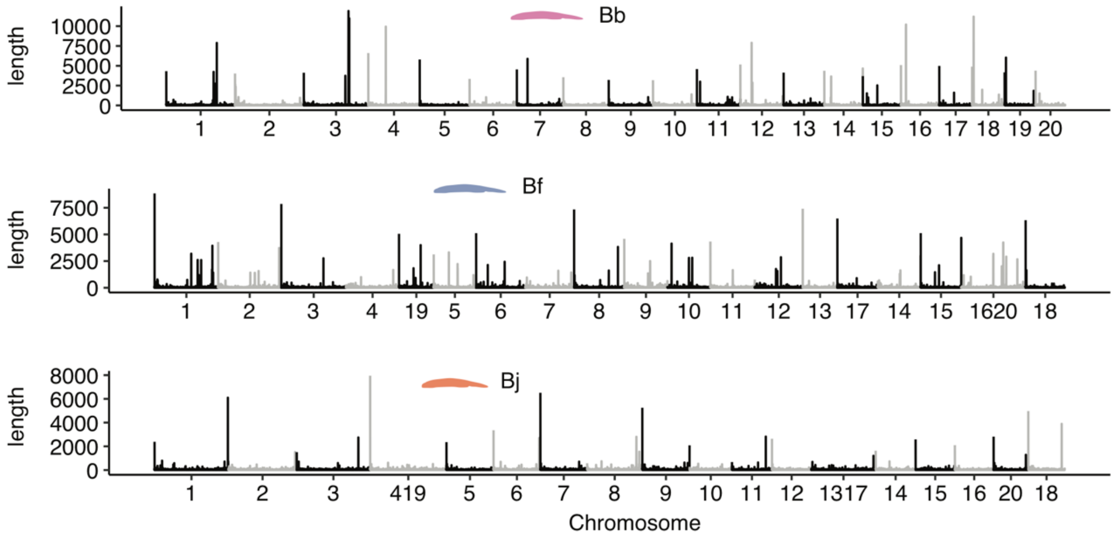


**Supplementary Fig. S10 The distribution of G4 sequences along the chromosomes**.
The G4 (G-quadruplex) sites were predicted and the total lengths were summed up in every 20 kb window along the chromosomes. The peaks of G4 are often present at the ends of chromosomes, but sometimes are also seen in the middle of chromosomes.

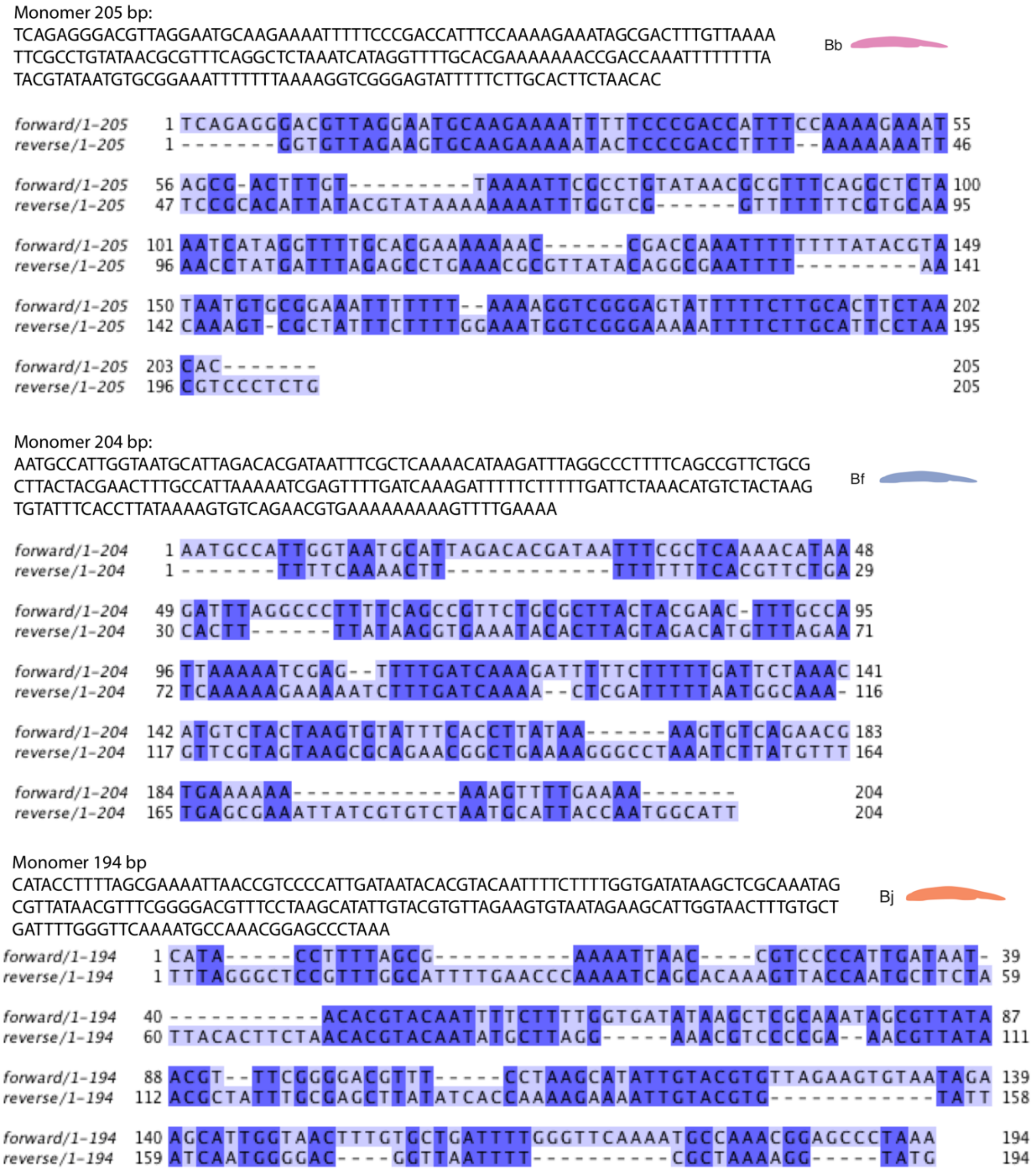


**Supplementary Fig. S11 The centromeric repeat monomer**.
The sequences of the centromeric repeat monomer are shown at the top of each panel. The monomers are not homologous among amphioxus species. For each monomer, the reversed complementary sequence (labelled as reverse) was generated and aligned with the original (labelled as forward) monomer. The dark blue highlights the identical nucleotides between the forward and reverse monomer.

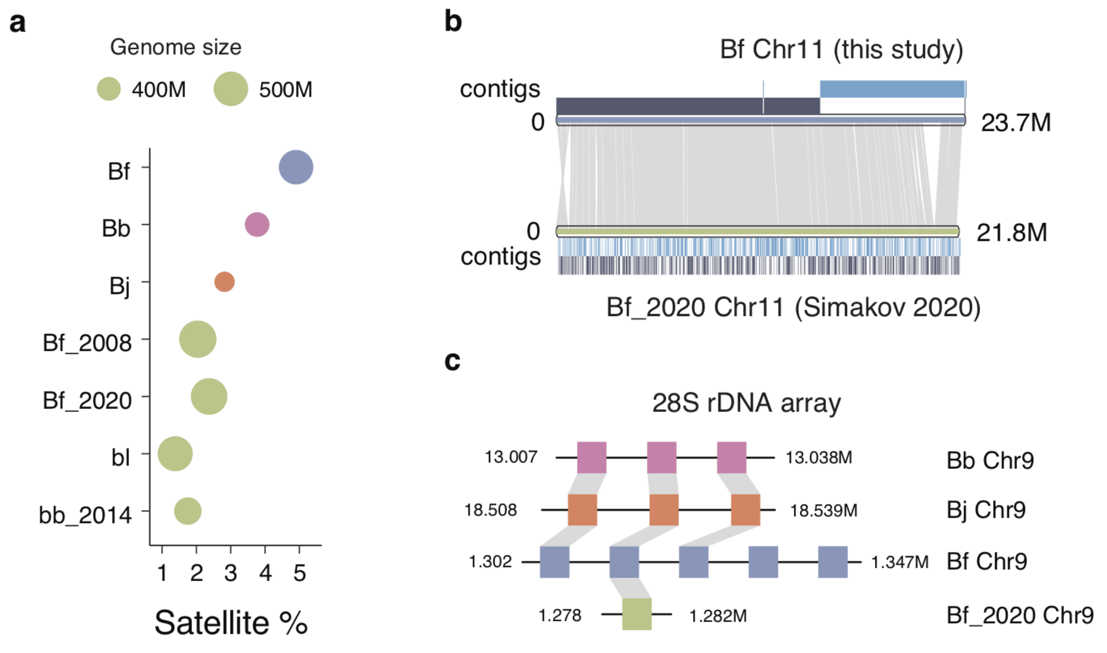


**Supplementary Fig. S12 Complex genomic regions of amphioxus species have been assembled by long-reads**.

**a)** In the long-read assemblies, the satellite sequences comprise a larger portion of the genome because more satellite sequences were assembled, compared to the previously published genomes. The circle size represents the genome size. The larger genomes have a higher portion of satellite repeats.

**b)** The genomic synteny of Chr11 between the long-read (Bf) and the published short-read (Bf_2020) assemblies. The grey bands represent syntenic blocks. The short-read assembly missed one large sequence fragment at the ~20Mb position. The dark and light blue blocks represent contigs. The Chr11 of long-read assembly is comprised of only 6 contigs, while that of the short-read assembly is comprised of thousands of contigs.

**c)** The gene synteny of a 28S rRNA gene array. There are 3, 3 and 5 tandem copies of 28S rRNA genes assembled in Bb, Bj and Bf long-read assemblies respectively, but only one copy was assembled in the short-read assembly of Bf.

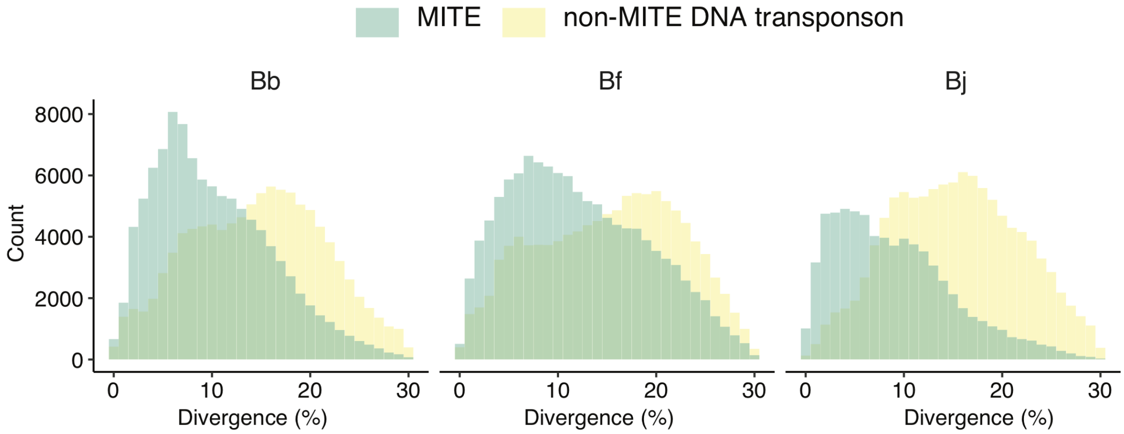


**Supplementary Fig. S13 The divergence of MITE in the amphioxus genomes**. MITE stands for Miniature Inverted-repeat Transposable Element which is a type of DNA transposons. The divergence of repeats (X-axis) were estimated by RepeatMasker. In all amphioxus species, MITEs have a lower divergence level from the consensus sequences, compared with other types of DNA transposon, suggesting more recent activities of MITEs.

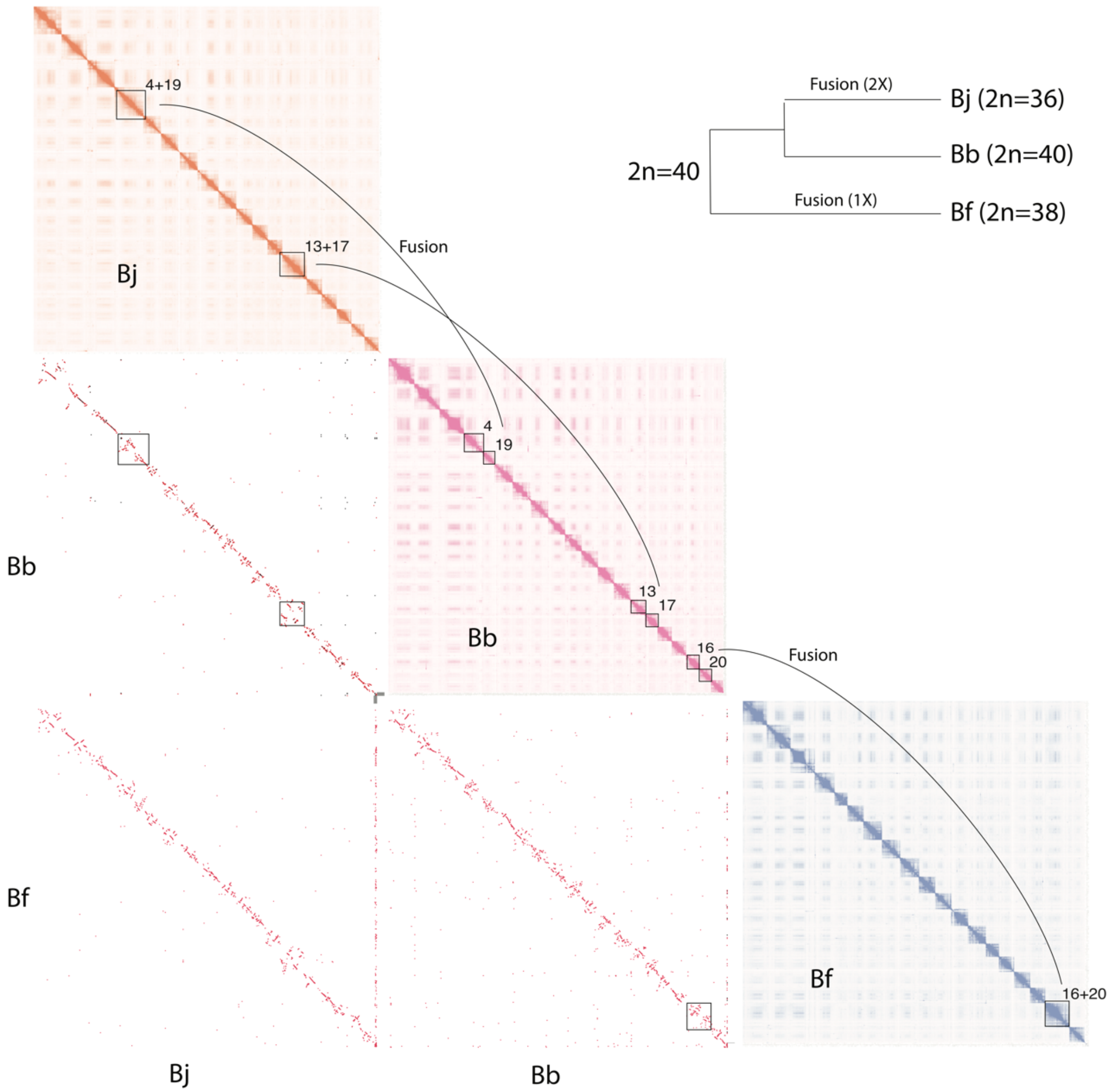


**Supplementary Fig. S14 Chromosomal fusions between amphioxus species**.
The Hi-C contact maps were visualized by JuiceBox. The genome synteny between each pair of species were shown underneath the Hi-C contact maps. The synteny dot-plots were visualised using the same methods described in Supplementary Fig. S11. The squares highlight the chromosomes that involve chromosome fusions. The numbers near the highlighted chromosomes in the Hi-C maps represent the chromosome numbers.

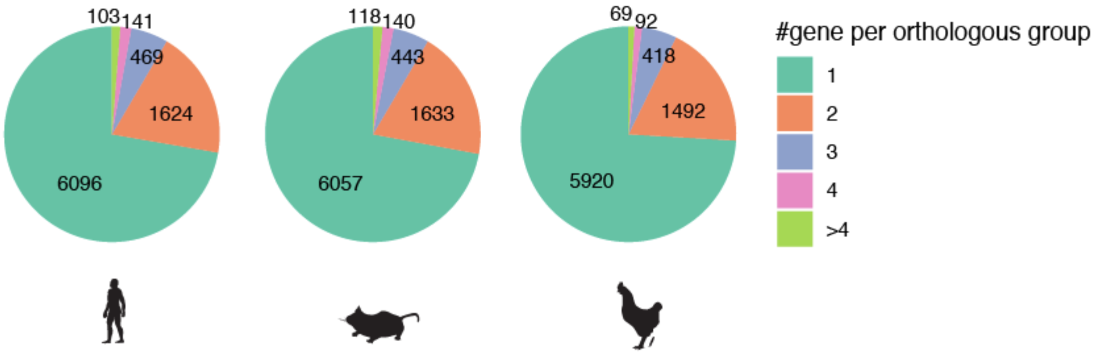


**Supplementary Fig. S15 Number of genes in each orthologous group**.
The statistics was directly retrieved from the OrthoFinder result. In most orthologous gene groups the genes are single copy. When the genes are multip-copy, the gene copy number mainly ranges from two to four.

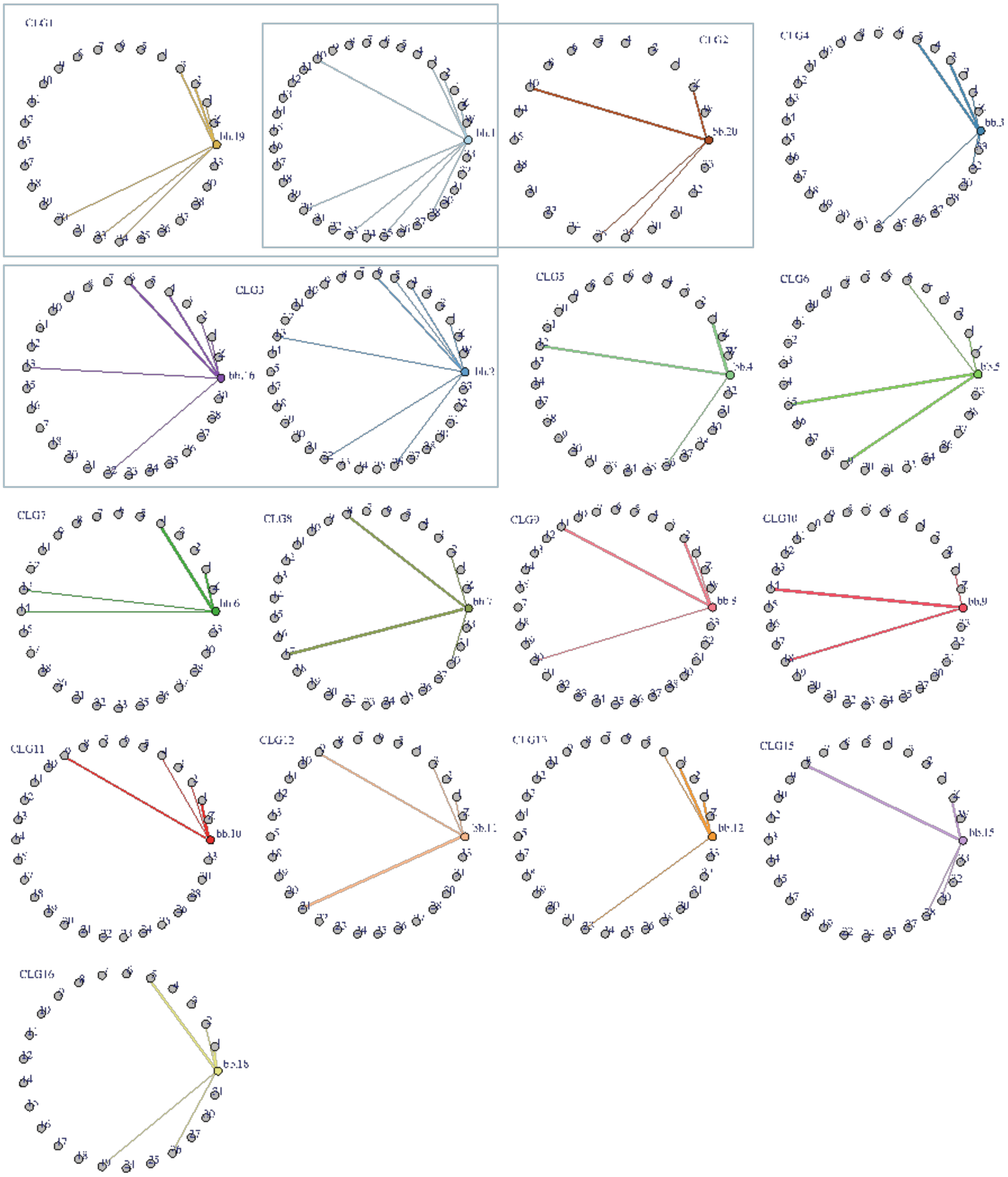


**Supplementary Fig. S16 Homology between chicken and Bb chromosomes**.
Each line connecting a Bb and a chicken chromosome represents the proportion of Bb genes (Bb%) of certain Bb chromosome that is homologous to the genes of the chicken chromosome. Only the lines with Bb% values larger than 4% are shown. One Bb chromosome usually connects with three or four chicken chromosomes. Each small panel represents one Bb chromosome that represents one CLG (except for CLG1, CLG2 and CLG14 ). CLG14 is shown in Fig. 2b. A part of Bb chr1 (bb.1) shares the chicken homology with chr19 while the other part shares with chr20 (The blue squares represent CLG1 and CLG2 respectively, both including a part of chr1).

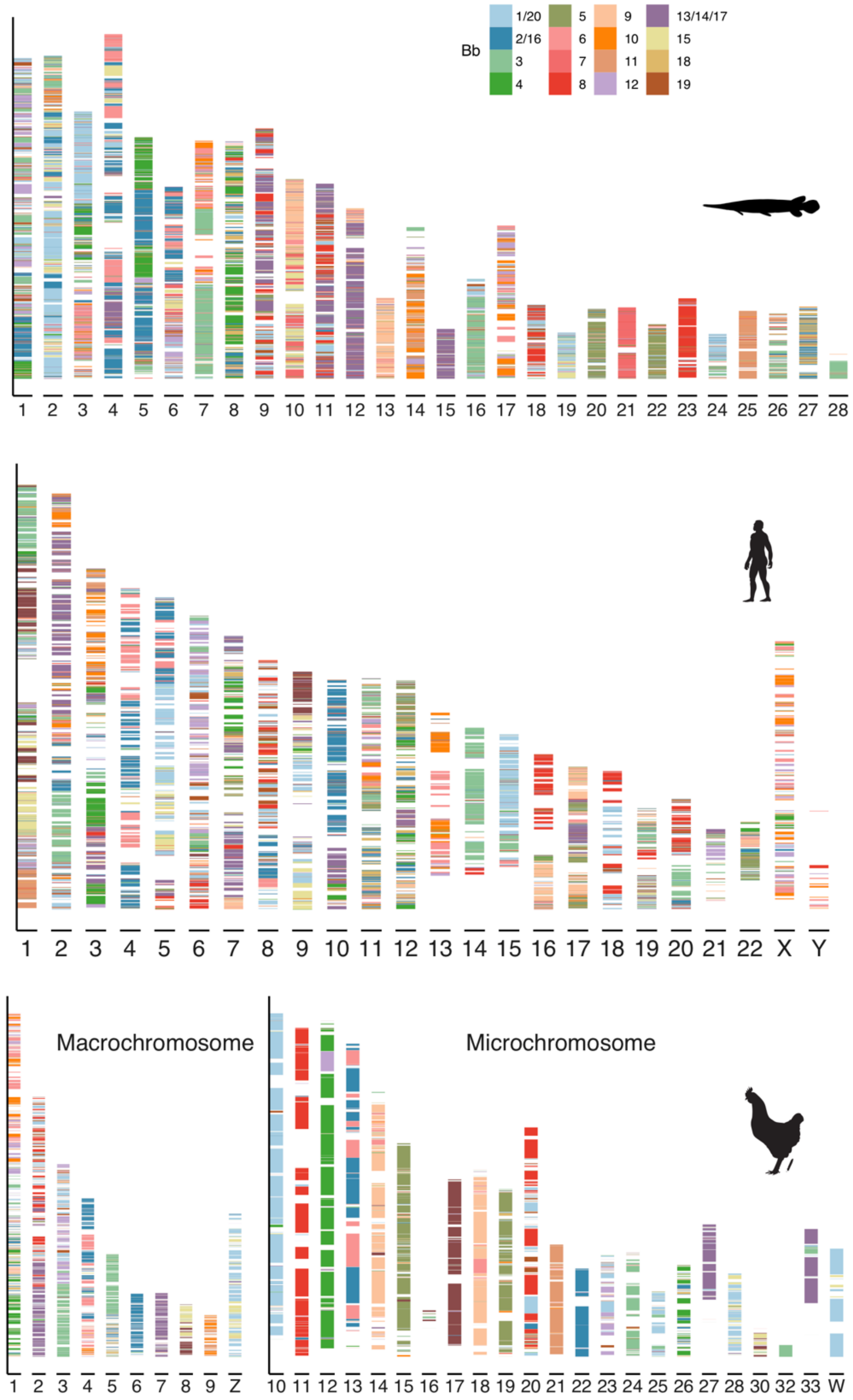


**Supplementary Fig. S17 The composition of human and spotted gar chromosomes by CLG**.

The colored bands show the Bb-chicken synteny blocks. A synteny block was built when two syntenic genes are not apart from each other for longer than 5 Mb. The Bb chromosomes were translated into CLGs according to the Bb-CLG relationship (Fig. 2c).

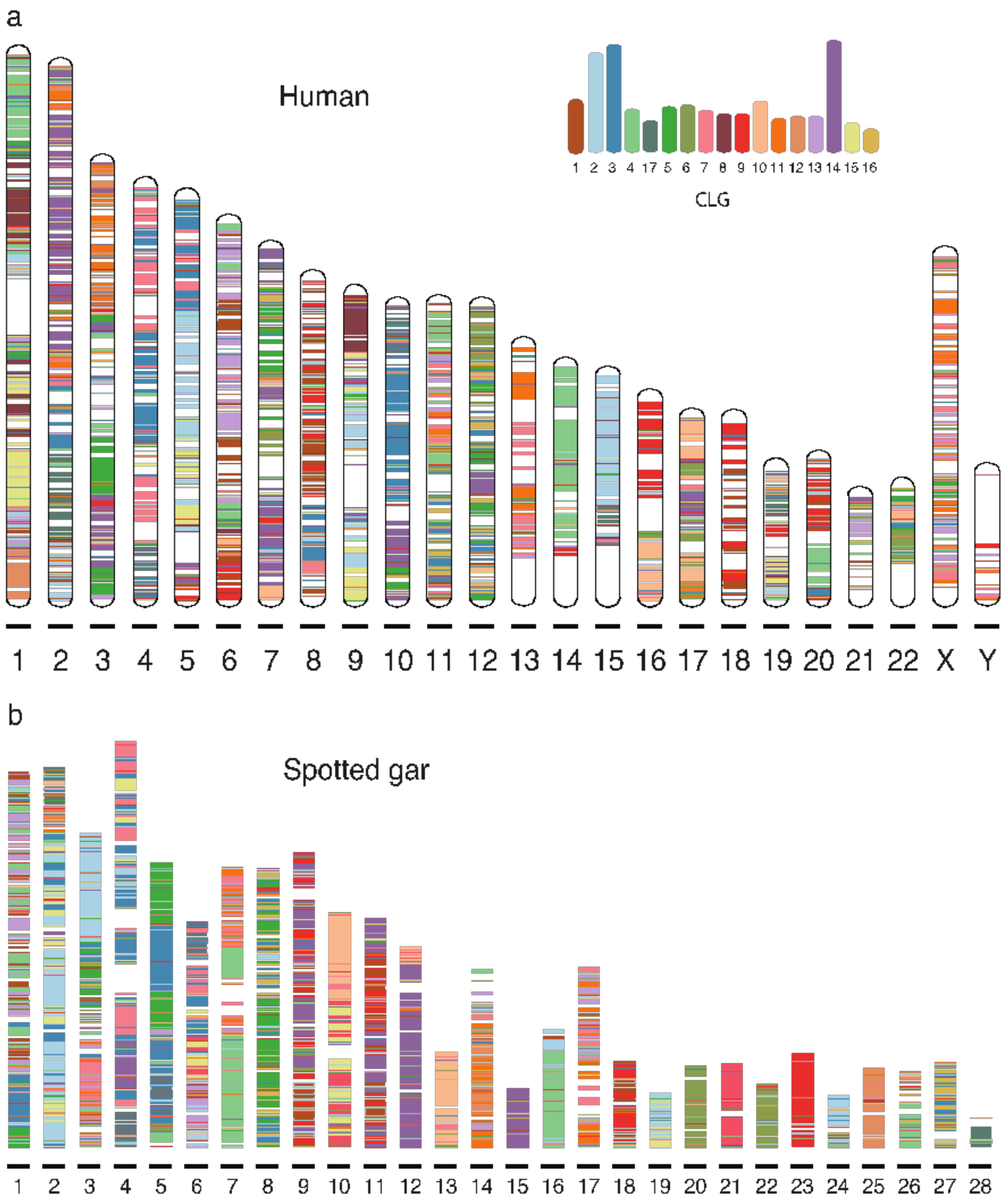


**Supplementary Fig. S18 The composition of human and spotted gar chromosomes by CLGs**.

Similar to Fig. 2d, we show the homologous sequences of CLGs across the chromosomes of human and spotted gar.

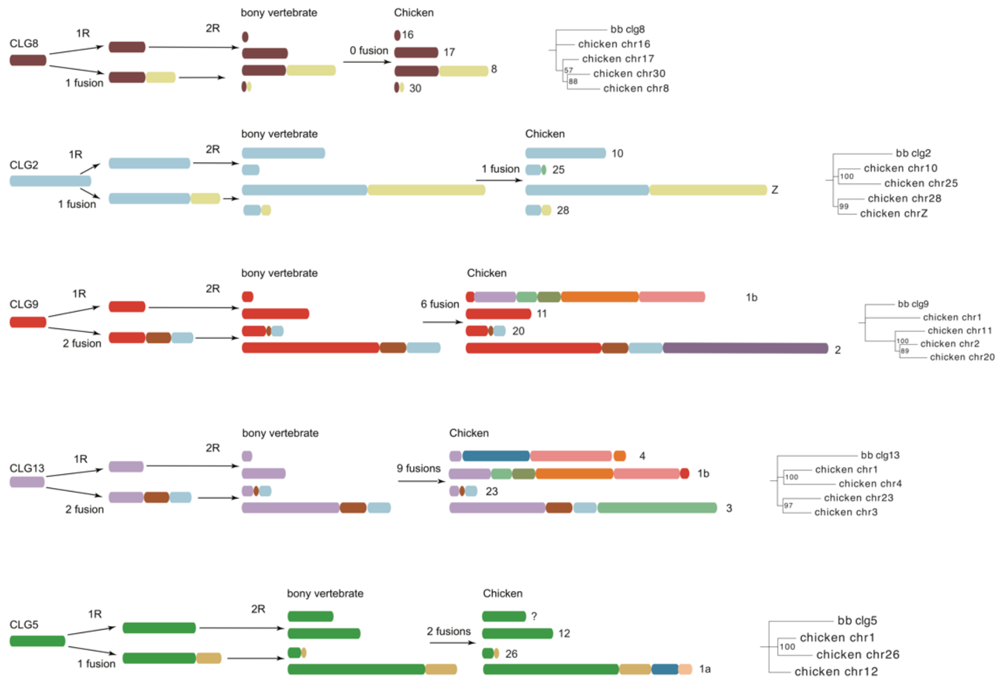


**Supplementary Fig. S19 Post-1R chromosomal fusions**.

The schematic plots show the evolutionary history of five CLGs, from the chordate ancestors to extant chicken chromosomes through 1R and 2R. A chromosome block in one color represents a CLG. For all five CLGs, one duplicated chromosome fused with another duplicate chromosome after 1R. Chicken chromosome numbers are labelled at the right end of the plot. In the right panels, the phylogeny was built using the ohnolog genes of chicken, using Bb as an outgroup. The bootstrapping values are shown at the nodes. The topology of the phylogeny supports the evolutionary relationships of ohno-chromosomes inferred based on post-1R fusions.

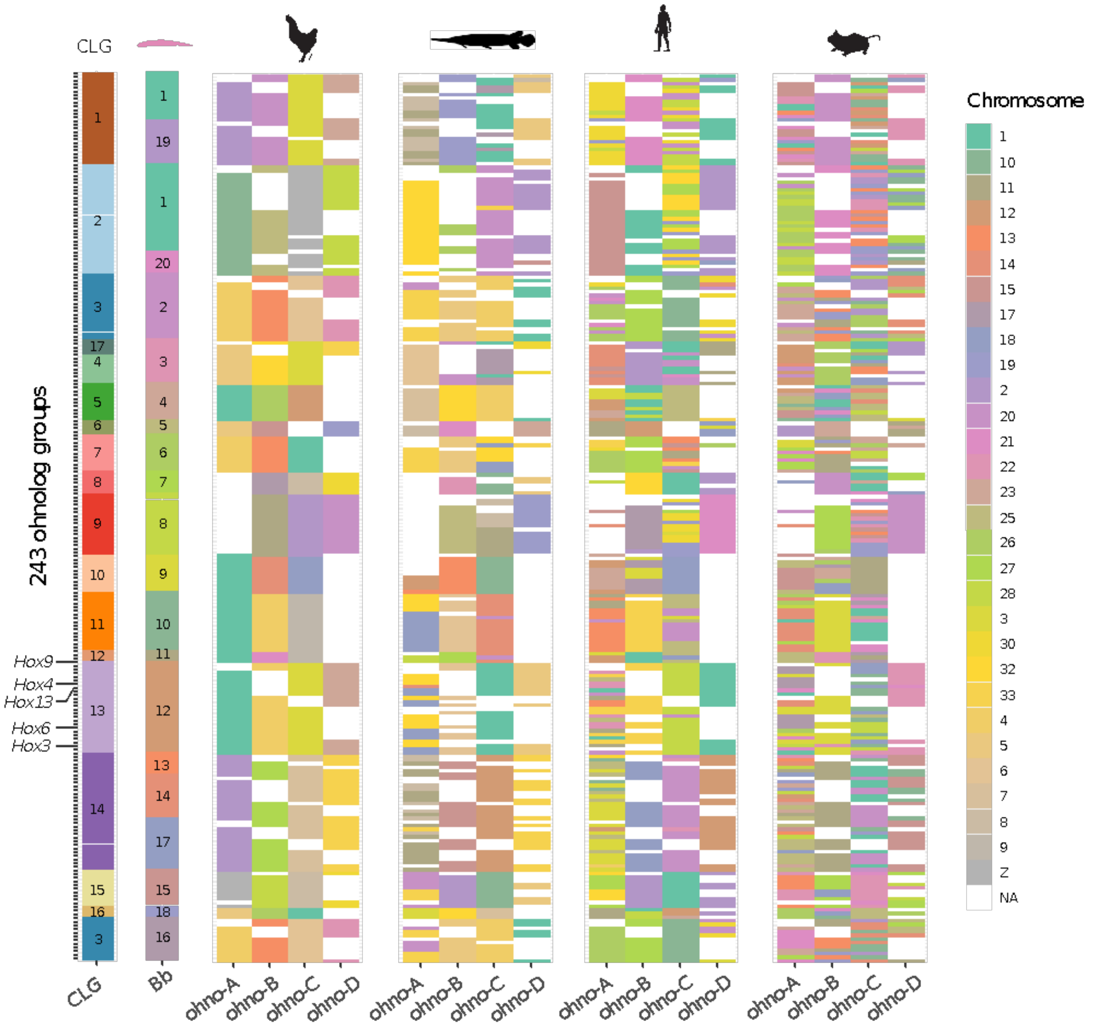


**Supplementary Fig. S20 Ohnolog groups used for dating the time of 1R and 2R**. There are 243 ohnolog groups identified that contain at least three vertebrate ohnologs. Each row represents one ohnolog group which contains one Bb gene, at least three chicken genes, 1-4 genes of spotted gar, human and mouse. The vertebrate genes of the same ohno-chromosome lineage (A/B/C/D) are homologous to each other. The IDs of CLGs and Bb chromosomes are labelled since they are clustered as large blocks, and the legend for vertebrate chromosomes was shown on the right. Five *Hox* genes are highlighted as examples.

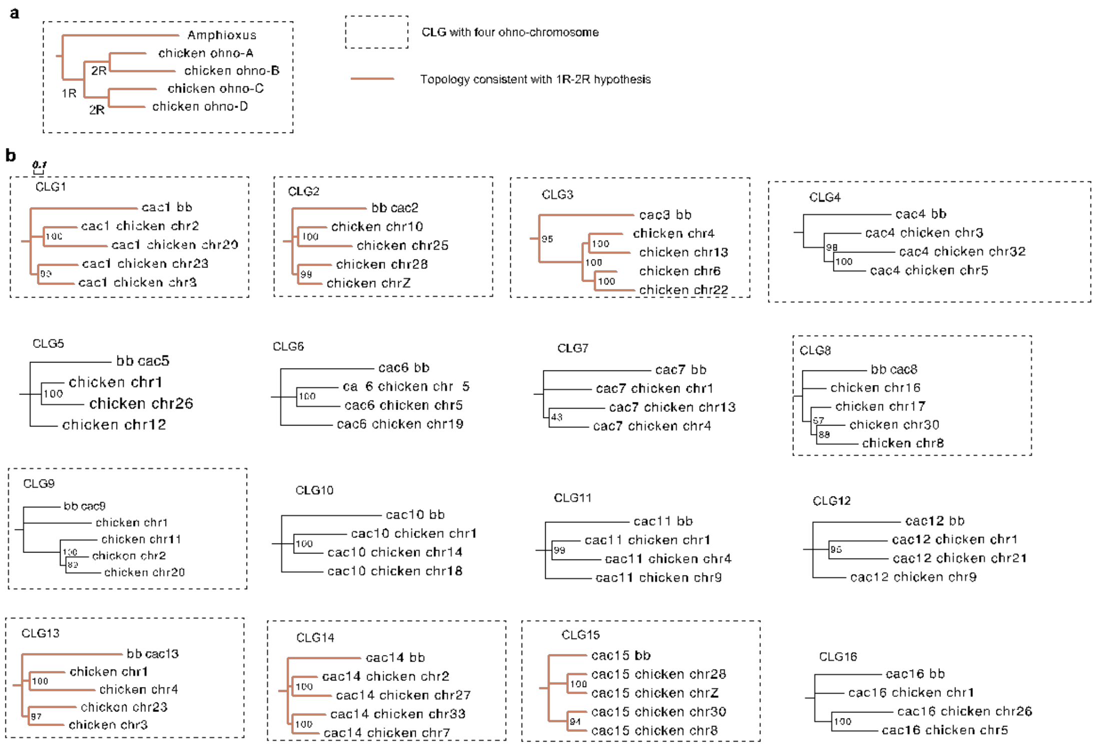


**Supplementary Fig. S21 Phylogeny of ohno-chromosomes built with ohnolog genes**. The phylogenetic trees of chicken ohno-chromosomes were built using the ohnologs gene residing on the same ohno-chromosomes. The list of ohnologs is listed after each ohno-chromosome branch. The phylogeny of the other five CLGs that experienced post-1R fusions in vertebrates is shown in Supplementary Fig. S15. The bootstrapping values are shown at the nodes.

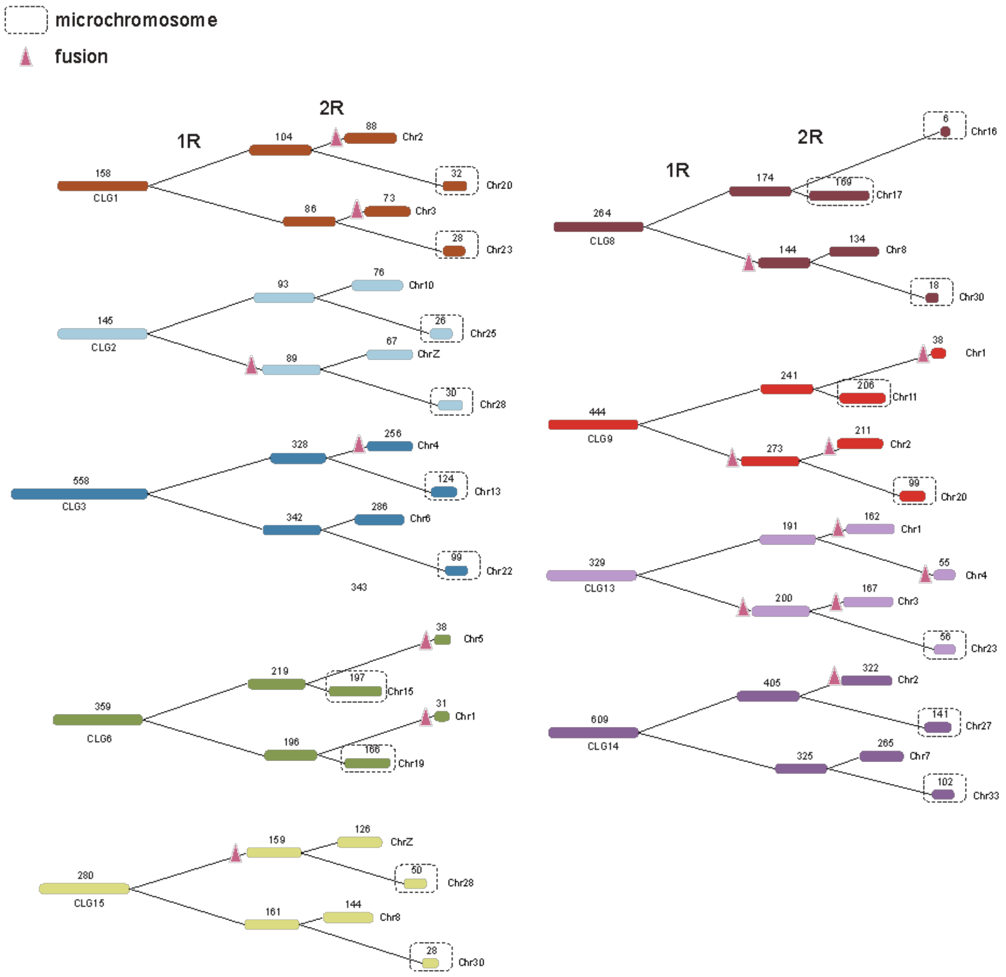


**Supplementary Fig. S22 Reconstruction of gene loss events following 1R and 2R**. The numbers above the CLGs are the original orthologous genes that have homologs in at least one descendant ohno-chromosomes. The numbers above the chicken and post-1R chromosomes show the extant and reconstructed gene numbers respectively. The branch length indicates the numbers of gene loss, *i.e.* a longer branch means more genes have become lost during or after WGDs in the branch. The pink triangles indicate fusion events along the branches. The microchromosomes are highlighted by dashed squares. In the branches leading to microchromosomes, no fusions have occurred, and the branches are longer (more gene loss).

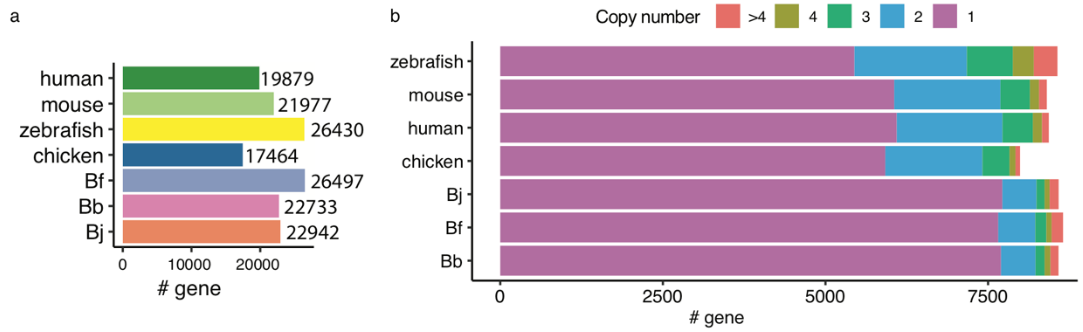


**Supplementary Fig. S23 Gene number of chordate genomes**.

**a)** The total number of protein-coding genes of four vertebrate species and three amphioxus species.

**b)** Statistics of gene counts in orthogroups. We included the orthologous gene groups in which at least one amphioxus gene and one vertebrate gene are present (*i.e.* the orthogroups are shared by amphioxus and vertebrates). Amphioxus has a larger proportion of single-copy genes, but also has a considerable amount of duplicated genes given the lack of WGDs.

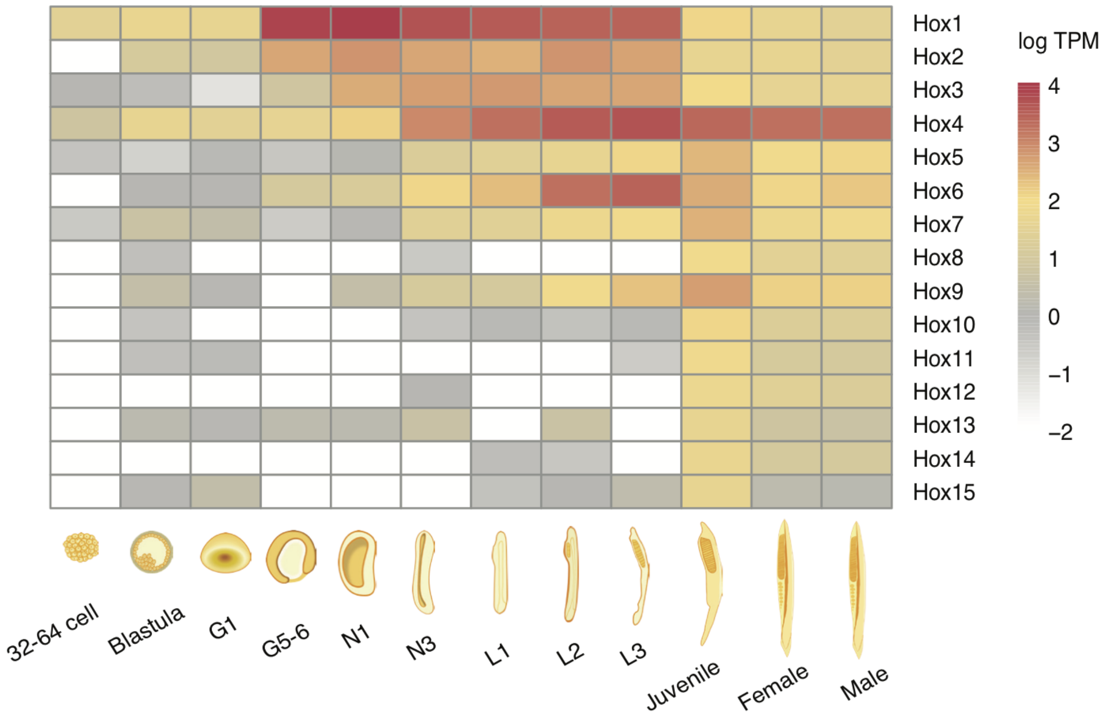


**Supplementary Fig. S24 The temporal expression of Hox genes**.
The expression profile of *Hox* genes of Bf across 11 developmental stages. In earlier stages only anterior Hox genes are expressed, and the posterior *Hox* genes are generally expressed in later stages.

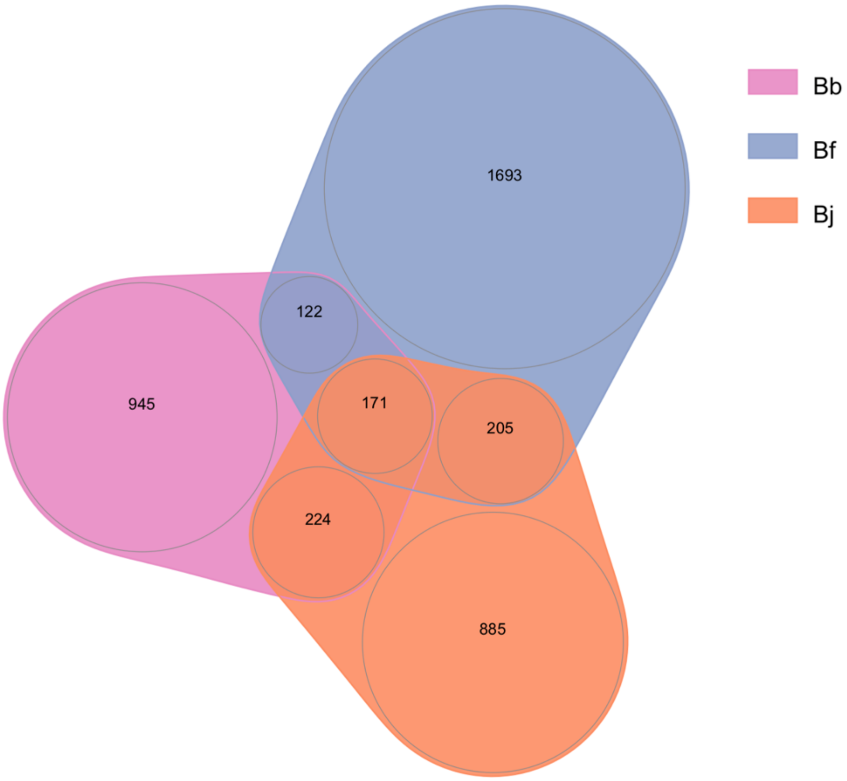


**Supplementary Fig. S25 The Venn diagram of genes involved in segmental duplications of three amphioxus species**.

For those genes whose coding regions are overlapped for 60% or more of the segmental duplication regions in three different amphioxus species, we identified the orthologous genes that are shared between or specific to the three species. We found that most of the genes are unique in one of the amphioxus species, suggesting the segmental duplications in amphioxus tend to be of more recent origin.

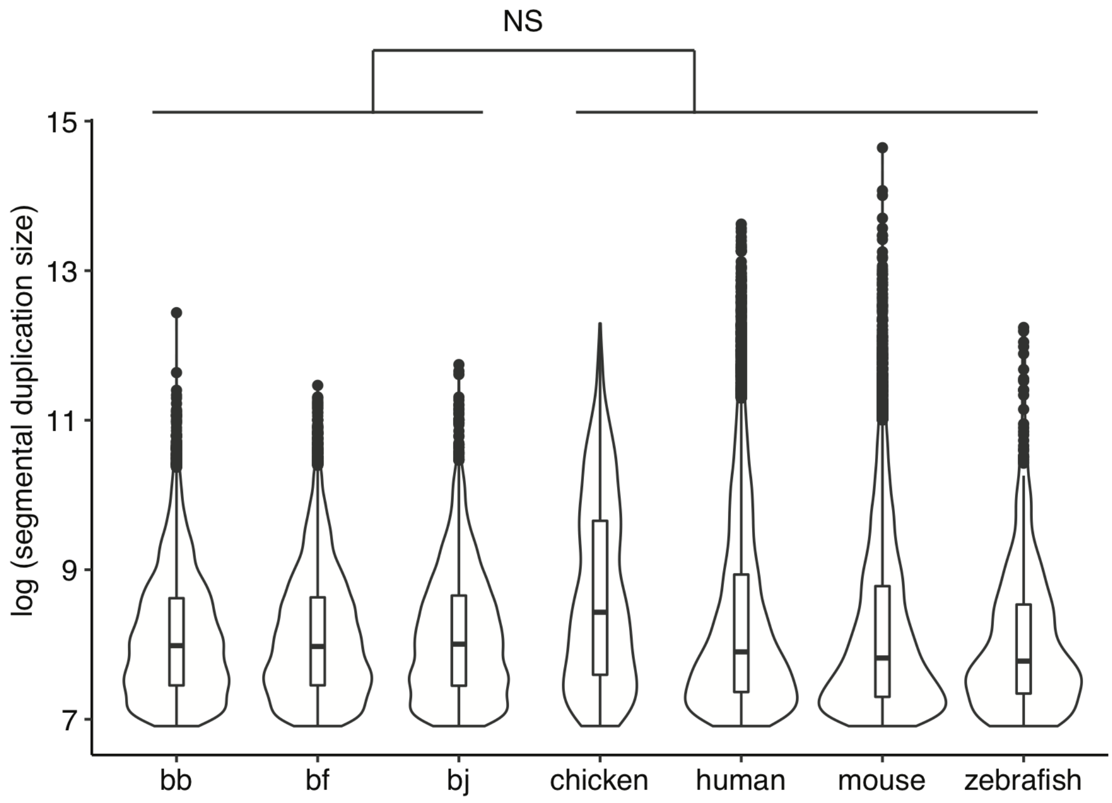


**Supplementary Fig. S26 The size of segmental duplications**.
The lengths of segmental duplications of the amphioxus genomes were compared with those from the vertebrate genomes. By definition all segmental duplications studied are larger than 1 kb. The lengths of segmental duplications of amphioxus are not significantly different from those of vertebrates (Wilcoxon test, $p > 0.05$).

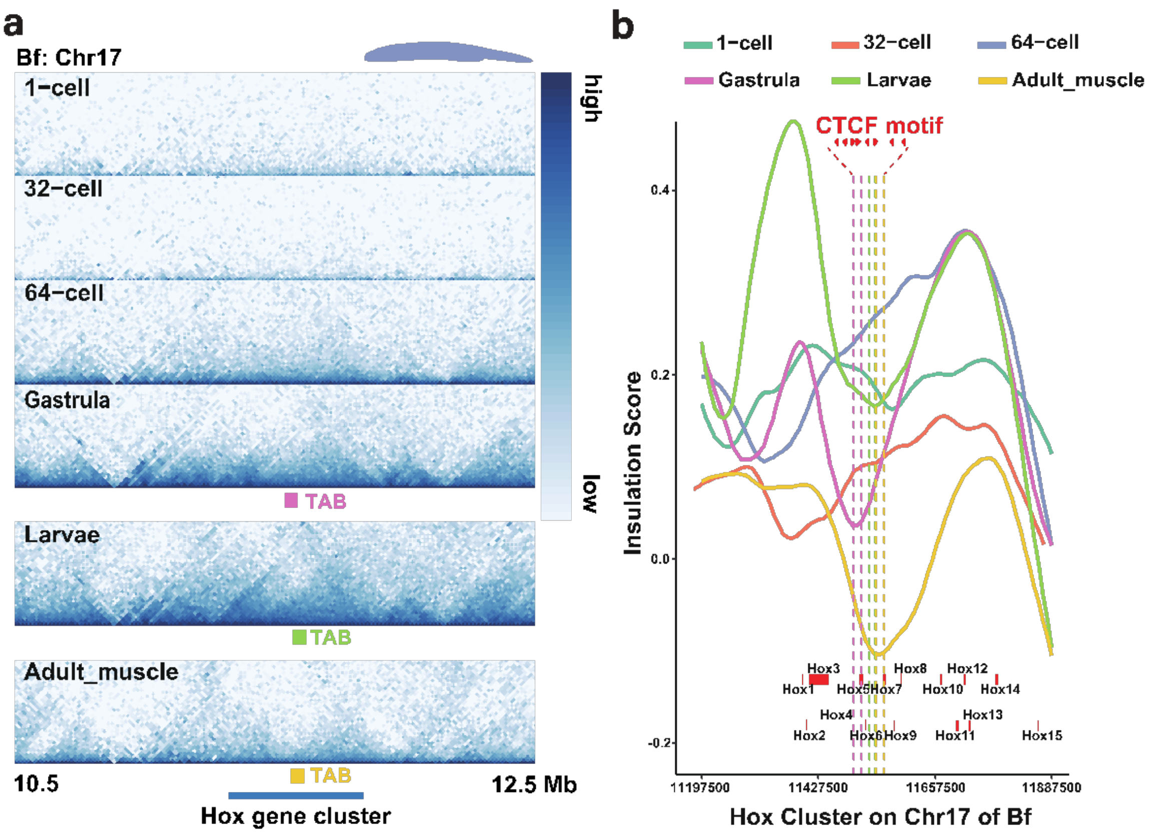


**Supplementary Fig. S27 TAD and insulation score at 15kb resolution within the amphioxus Hox gene cluster.**

a) The interaction heat map for high-order chromatin structures of chromosome 17(10500000-12500000) showing the TABs within *Hox* cluster is shown for 1-cell, 32-cell, 64-cell, gastrula, larvae and adult(muscle tissues) stages of Bf at 15-kb resolution. The dots under heat maps of Gastrula, Larvae and Adult_muscle indicate the TABs of these three developmental stages.

b) The horizontal solid lines indicate the distribution of insulation scores along the *Hox* gene cluster, the smaller the insulation score, the stronger the TABs. The vertical dotted lines represent the TABs of gastrula (pink), larvae (green) and adult (yellow) stages of amphioxus. The length and location of the *Hox* genes are marked with thick red lines. The TAB at the *Hox* gene cluster is shifted between different developmental stages, but between the regions of *Hox5* and *Hox7*.

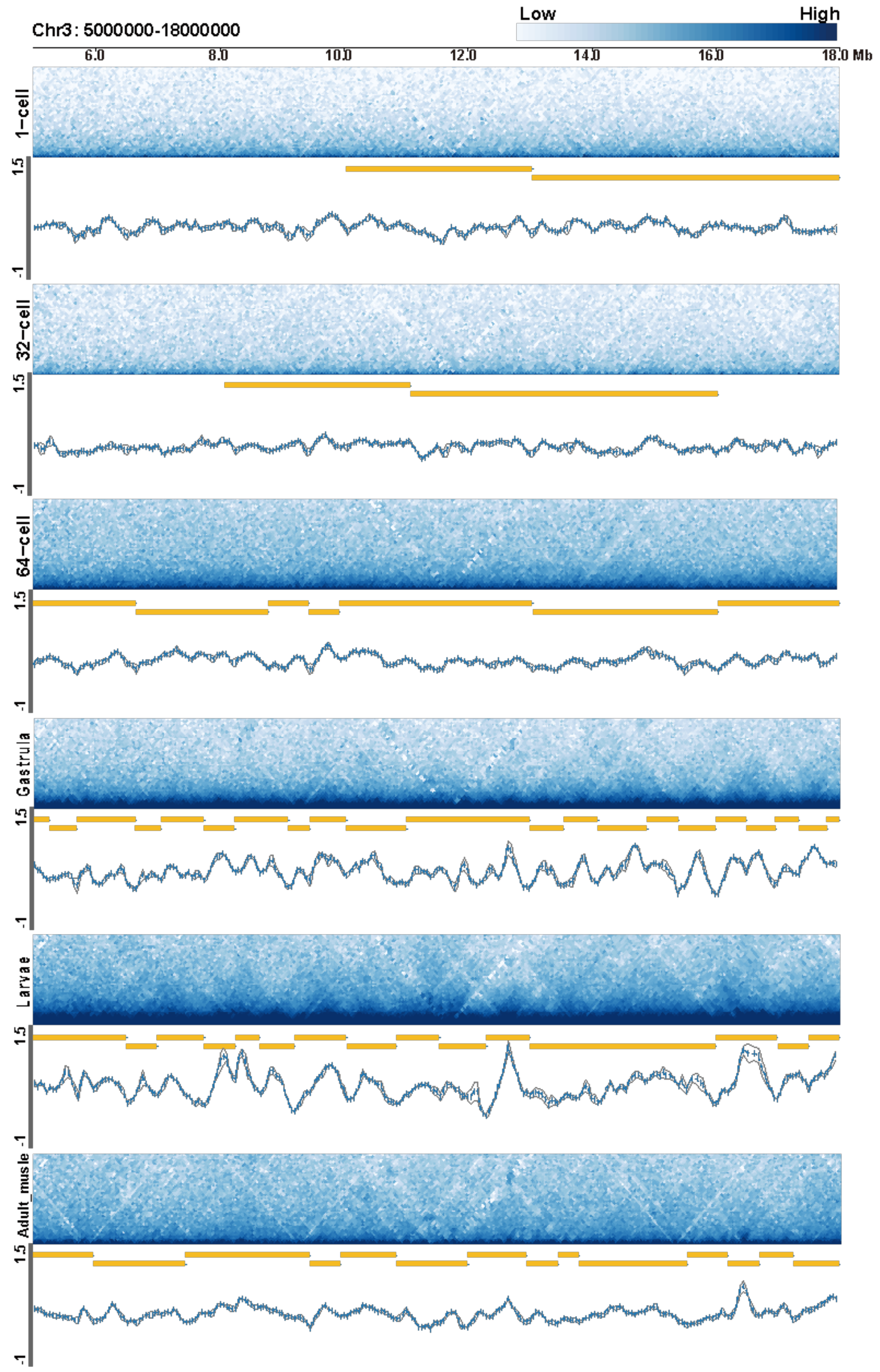


**Supplementary Fig. S28 Three-dimensional chromatin structures during embryonic development of amphioxus**.

The interaction heat map for high-order chromatin structures of chromosome 3(5M-18M), as an example, is shown for 1-cell, 32-cell, 64-cell, gastrula, larvae and adult(muscle tissues) stages of Bf at 50-kb resolution. The lines indicate the insulation score of each development stages.

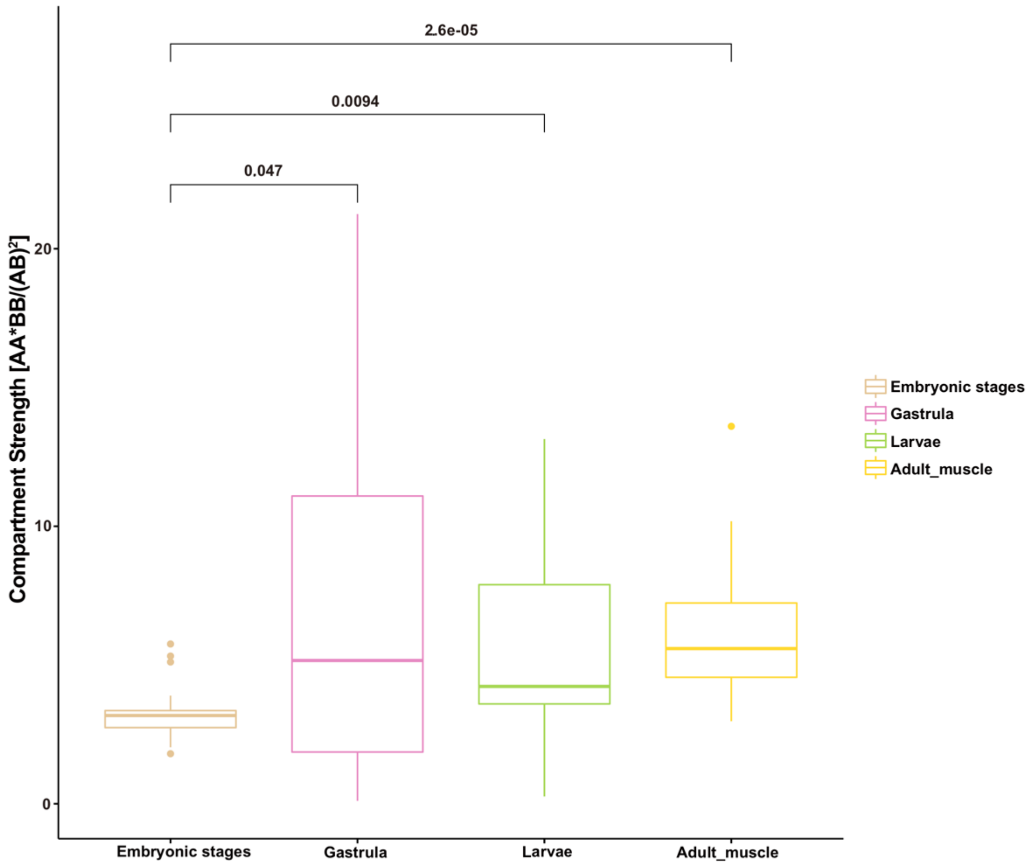


**Supplementary Fig. S29 A/B compartment strength between different developmental stages**

Compartment strengths were calculated as $AA*BB/AB^2$ for each chromosome. The value in embryonic stages is the average value of compartments strength identified in 1-cell, 32-cell and 64-cell.

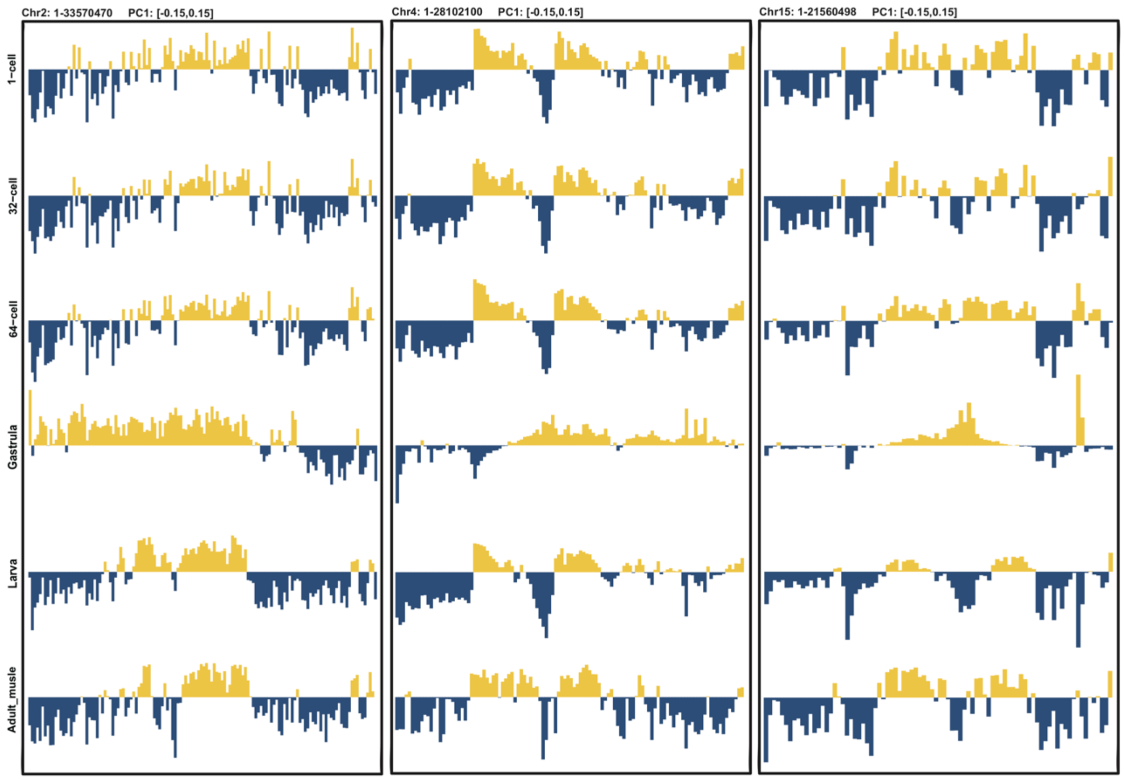


**Supplementary Fig. S30 Eigenvector 1 value tracks for amphioxus chromosomes at 250kb resolution at six different developmental stages**.

Eigenvector 1 values (PC1, 250kb resolution) across the amphioxus chromosome 2, chromosome 4 and chromosome 15, representing A (yellow) and B (blue) compartments. On some chromosomes, e.g., chr2, compartment strength is significantly reduced at the gastrula stage.

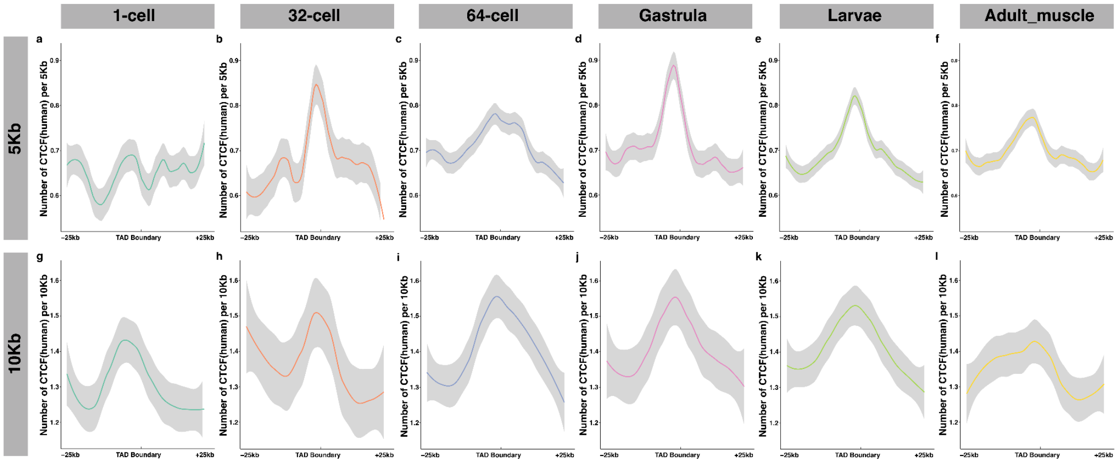


**Supplementary Fig. S31 Enrichment of putative CTCF-binding sites at TAD boundaries in the amphioxus genome**.

a-f) Putative CTCF binding site density was calculated in 5kb windows overlapping each other 50% along the up- and down-stream (25kb) regions of amphioxus TAD boundaries. For each window, the mean value of the CTCF density was used to generate the plots. Panel a-f represent 1-cell, 32cell, 64-cell, gastrula, larvae and adult muscle respectively.

g-l) The same as panel a-f but in 10kb resolution.

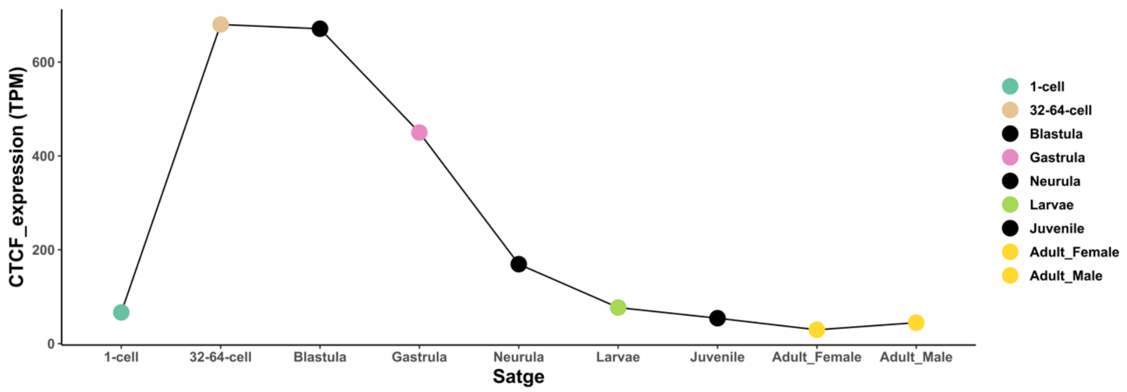


**Supplementary Fig. S32 Expression of CTCF in Bf during developmental stages**. The expression of the CTCF gene is increased at zygotic genome activation (ZGA) around the 64-cell stage.

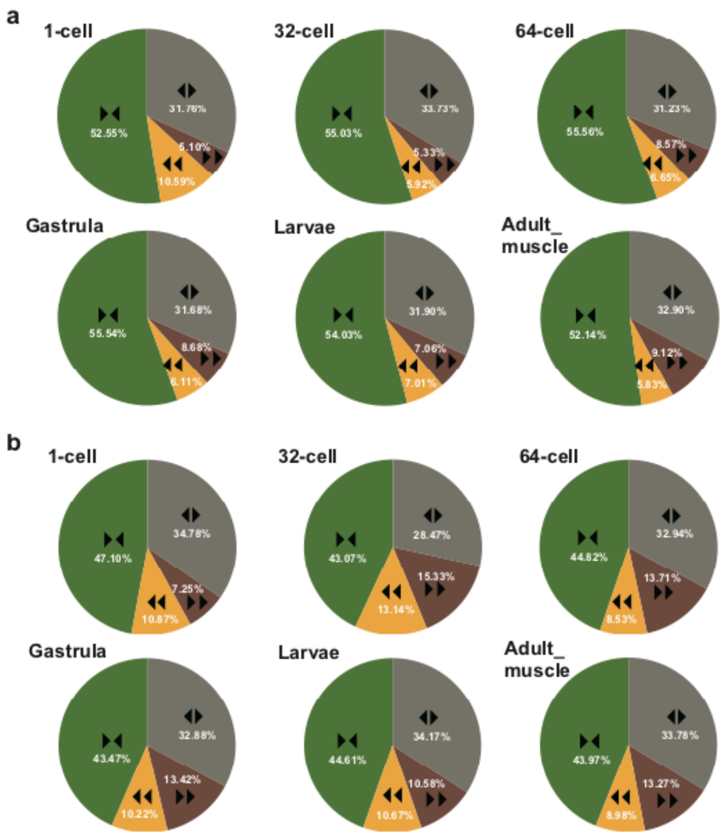


**Supplementary Fig. S33 The orientation of CTCF-binding sites pairs in amphioxus TABs.**

a) Pie chart shows four types of orientation (arrowheads and different colors, green color for the convergent CTCF site pairs) for the paired CTCF sites in the TABs. The TABs are identified at the 5kb resolution and extended 5kb up- and down-stream, respectively. The convergent CTCF-binding site pairs are the major type(>52%).

b) The same as a, but in 10kb resolution. The ratio of convergent CTCF-binding site pairs reduced to about 45%.

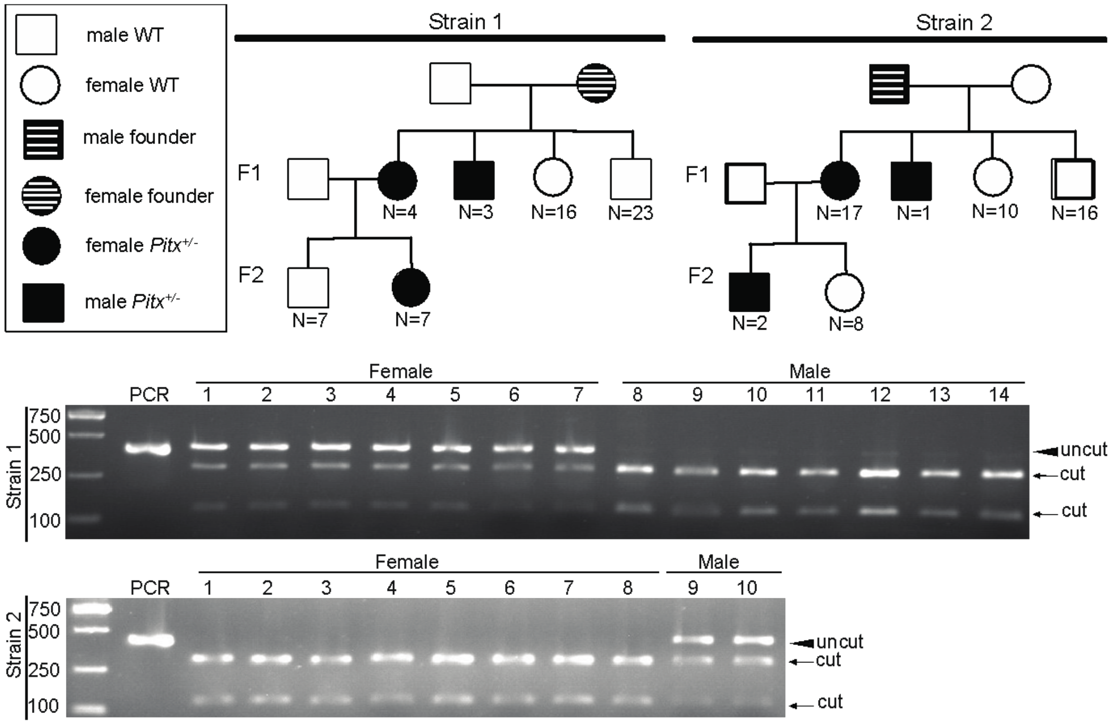


**Supplementary Fig. S34 Genetic evidence for the ZW sex chromosomes of *B. floridae***

Pedigree and genotyping of *Pitx* mutant strains. The mutation types carried in *Pitx*$^{+/-}$ heterozygotes of this figure include frame shift and non-frame shift mutations. The animal numbers analyzed are noted. Strain 1: a female founder injected with TALEN pair Pitx-Fw3/Rv3 (primer Fw3: 5'-GCAACCGTTCGACGAC-3' and Rv3 5'-TGTAGGCCGGCGAGTA-3') was crossed with a WT male. One F1 female *Pitx*$^{+/-}$ heterozygote carrying mutation (-14 bp) was crossed a WT male.

Strain 2: a male founder injected was crossed with a WT female. One F1 female *Pitx*$^{+/-}$ heterozygote carrying mutation (-7 bp, +18 bp) was crossed with a WT male. Genotyping analysis of F2 progenies from both strain 1 and 2 is present. Amplicons from were digested with *Tat*I enzyme and separated on 2% agarose gel to identify their genotypes.

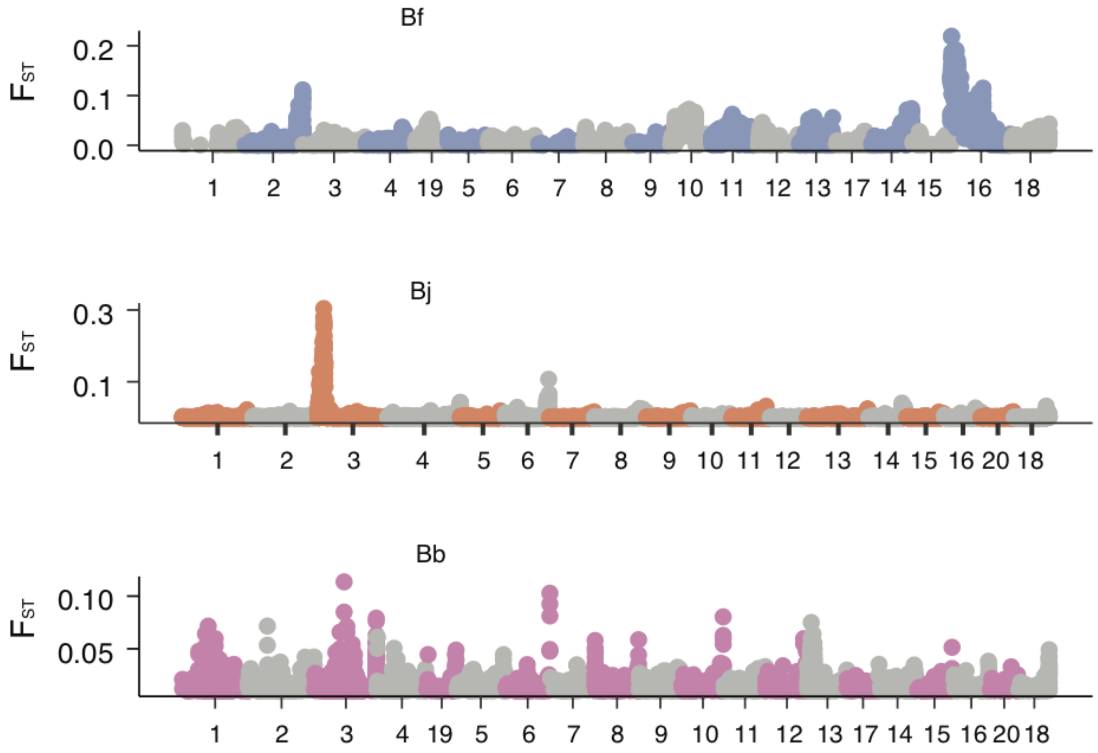


**Supplementary Fig. S35 The $F_{ST}$ statistics between male and female populations**. Only biallelic SNPs were considered for $F_{ST}$ calculation. The $F_{ST}$ values were estimated in 10 kb windows for Bj and Bf, but 5 kb for Bb because of the smaller size of the non-recombining region (~1.5 kb) in Bb. The highest peaks of $F_{ST}$ appear at Chr16, Chr3 and Chr3 of Bf, Bj, and Bb respectively. The number of individuals re-sequenced can be found in the Supplementary Table S6.

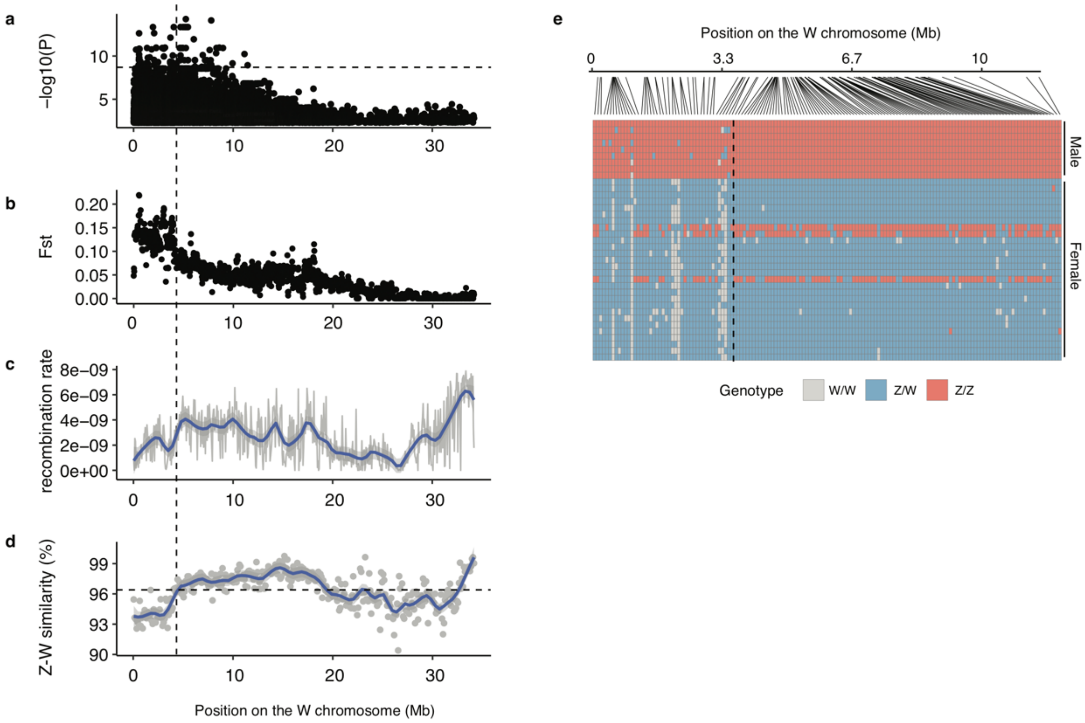


**Supplementary Fig. S36 The sex-linked region in Bf.**

**a)** A zoom-in view of Fig. 5a on the Chr16 (ChrW). **b)** A zoom-in view of Fig. S34 on the Chr16 chromosome. **c)** This panel was retrieve from Fig. S7 for Bf Chr16. **d)** We used Lastz to align the ChrZ and ChrW sequences, and calculated the sequence similarity in 100 kb windows (grey dot). The blue line shows the smoothed mean values. The horizontal dashed line shows the mean sequence similarity between the Bf and Bf_2 genomes. The vertical dashed line shows the boundary of the first evolutionary stratum. **e)** The position and genotype of the female-associated SNPs. Each column in the lower panel represents a SNP site and each raw represents an individual. The vertical dashed line shows the boundary of the first evolutionary stratum

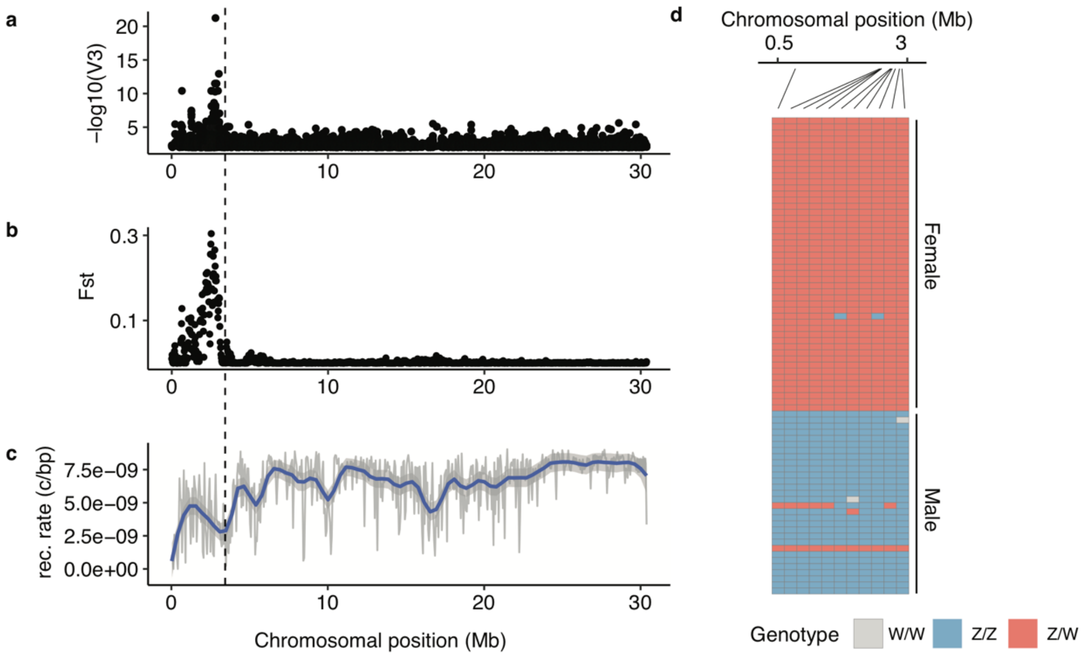


**Supplementary Fig. S37 The candidate sex determining region of Bj**.
**a)** A zoom-in view of Fig. 5b on the Chr3 (ChrZ). **b)** A zoom-in view of Supplementary Fig. S34 on the Chr3 chromosome of Bj. **c)** This panel (recombination rate) was retrieved from Supplementary Fig. S7 for Bj Chr3. The vertical dashed line indicates the boundary of the sex-linked region. **d)** The position and genotype of the female-associated SNPs. Each column in the lower panel represents a SNP site and each row represents one individual.

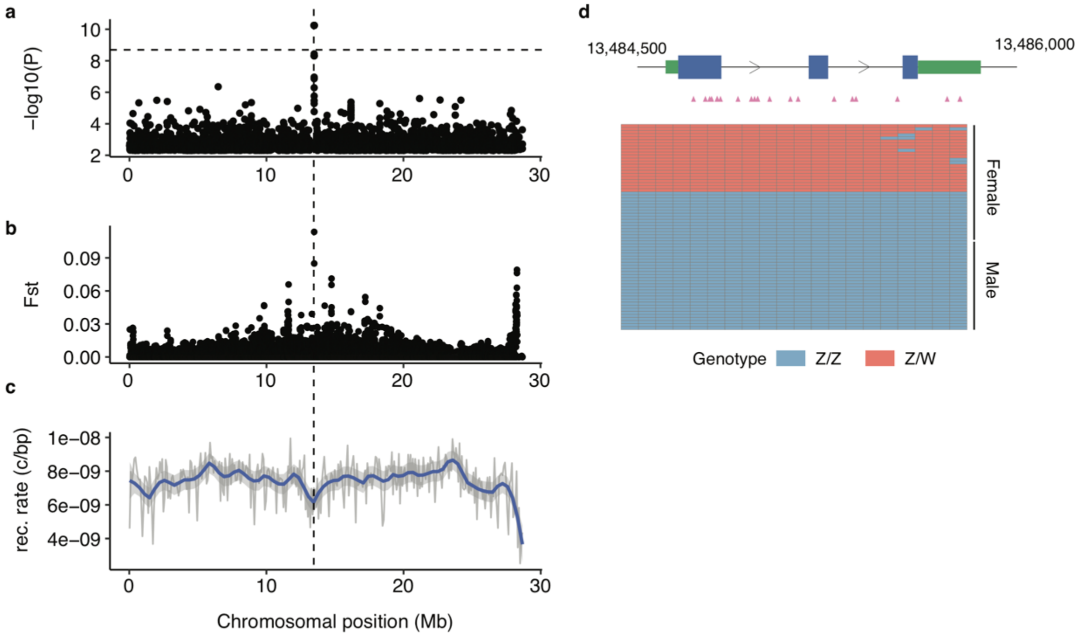


**Supplementary Fig. S38 The candidate sex determining region of Bb**.
**a)** A zoom-in view of Fig. 5c on the Chr3 (ChrZ). **b)** A zoom-in view of Supplementary Fig. S34 on the Chr3 chromosome of Bb. **c)** This panel (recombination rate) was retrieved from Supplementary Fig. S7 for Bb Chr3. The vertical dashed line indicates the position of the Bb sex-linked locus. **d)** The sex-linked variants (pink triangles) are all located in the gene body of the Bb candidate sex-determining gene. Each column in the lower panel represents a SNP site and each row represents an individual. The SNP sites are ordered according to their position in the genome.

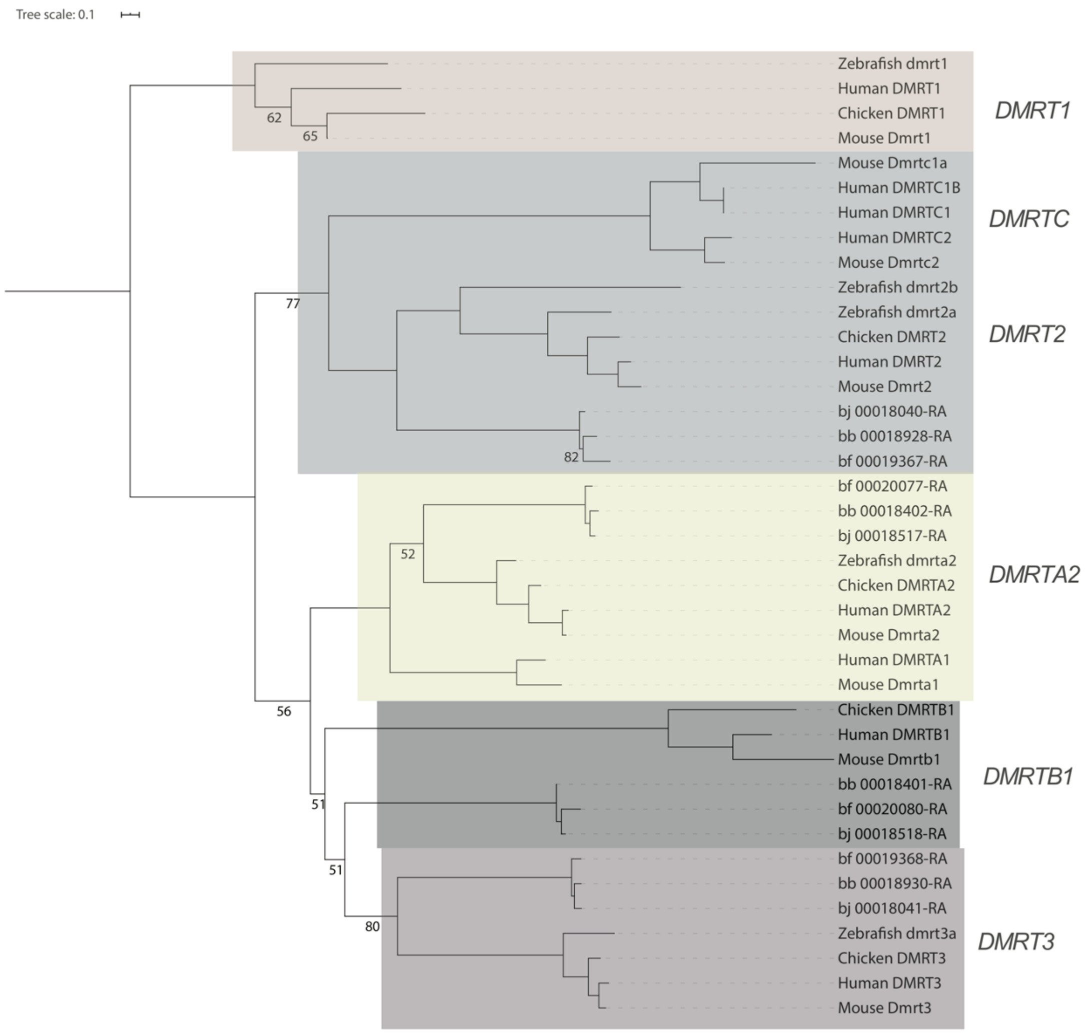


**Supplementary Fig. S39 The phylogeny of DMRT family genes across chordates**. The phylogeny was built from the coding sequences of the DMRT family genes of Bj, Bb, Bf, human, mouse, chicken and zebrafish. The root of DMRT family phylogeny was decided according to Mawaribuchi *et al.* 2019. The same families were highlighted in the same colored background. Bootstrapping values higher than 90 were not labelled at the nodes. Amphioxus lacks the *DMRT1* gene.

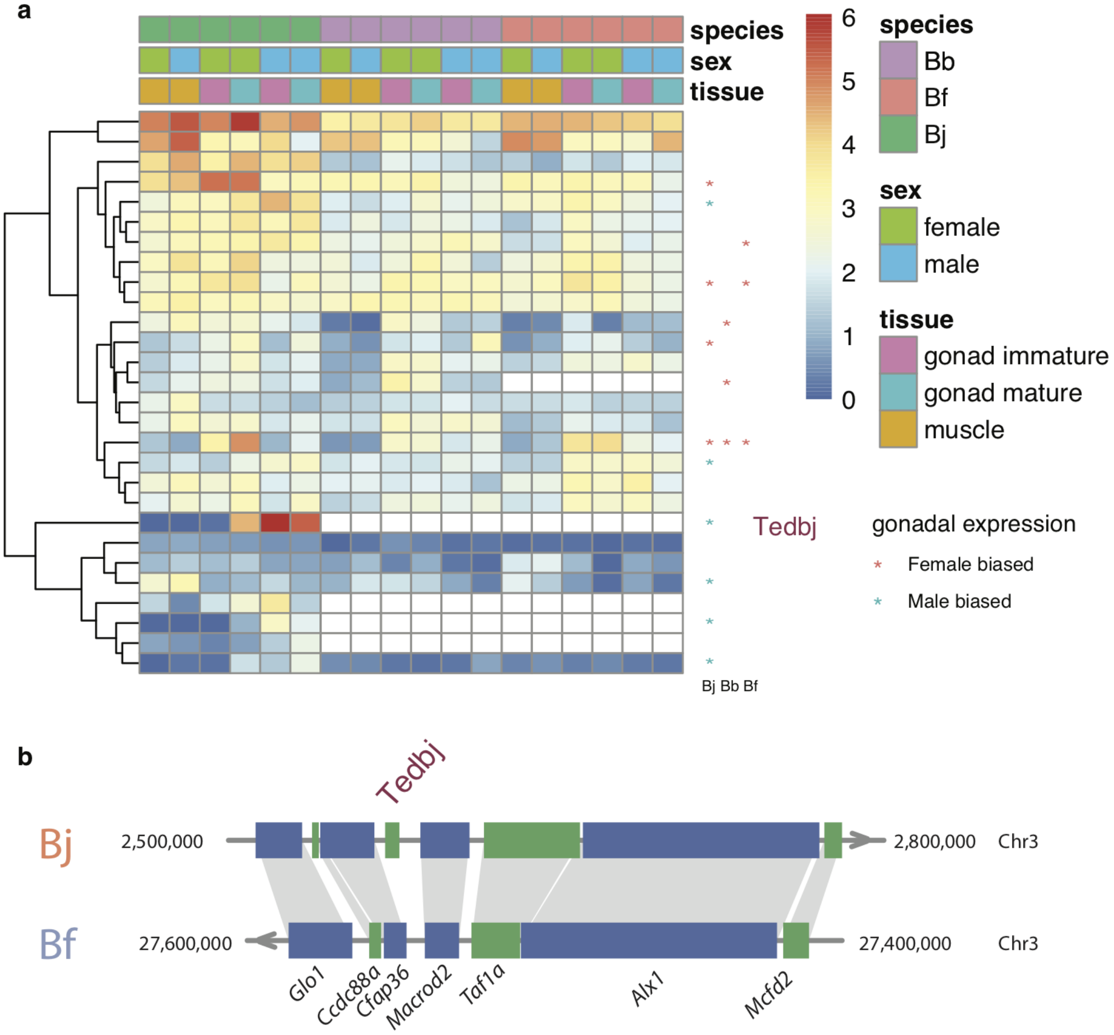


**Supplementary Fig. S40 The expression profile of sex-linked genes in Bj**.

**a)** The heatmap shows the expression levels of the sex-linked genes of Bj stratum I in immature and mature gonads, as well as muscle, in three amphioxuses. The expression levels were log1p transformed. The blank tiles in the Bb and Bj panels represent those genes whose orthologs are absent in certain species. The asterisks at the end of the rows indicate whether the genes show differential expression between the immature gonads of males and females in Bj, Bb and Bf. *Tedbj* was highlighted which has a strong testis-biased expression.

**b)** Gene synteny around *Tedbj* between Bj and Bf. *Tedbj* is found to be specific to Bj which is also lacking in Bb (not shown).

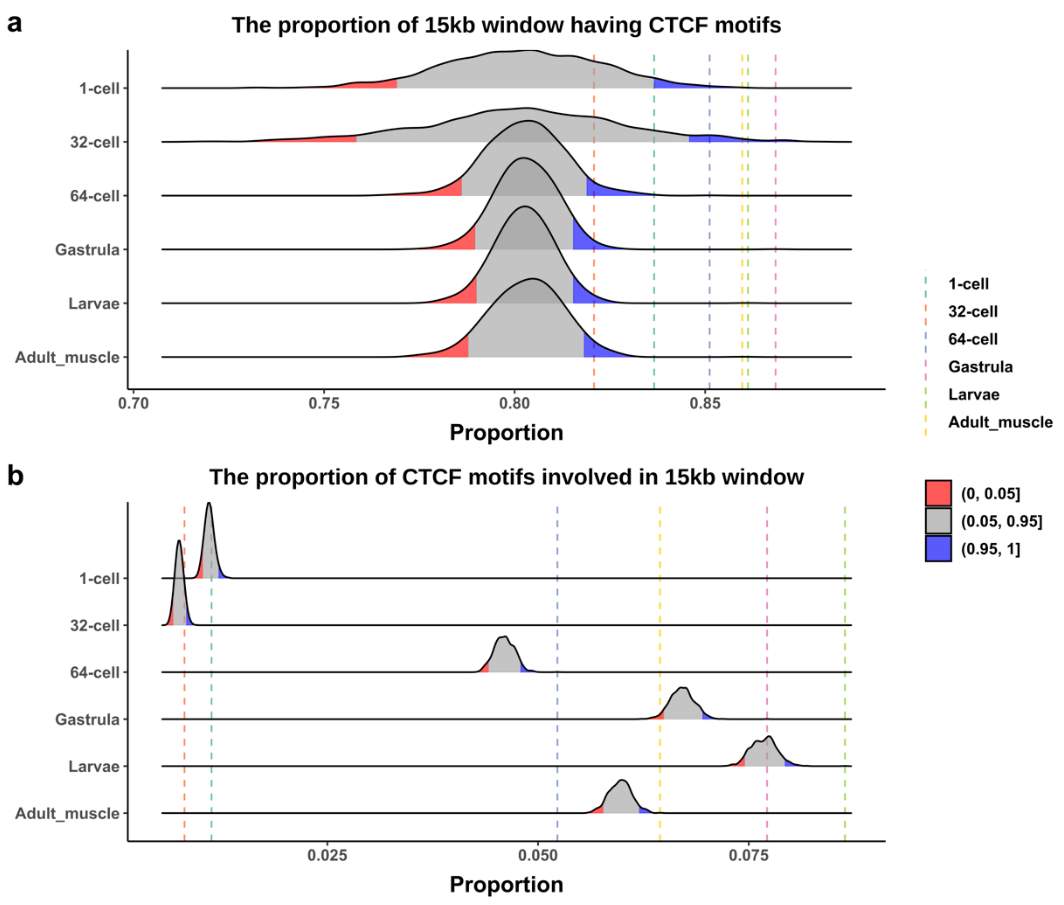


**Supplementary Fig. S41 Significance test of CTCF enrichment at TABs.**

**a)** For each developmental stage, we randomly selected 15kb windows with the same number of TABs across the genome and calculated the proportion of 15kb windows having CTCF motifs. We repeated this step for 1000 times and obtained the density of all numbers of proportions. The red part and blue part showed the lowest 5% and highest 5% proportions, respectively. The proportion of TABs having CTCF motifs of each stage was indicated by vertical dotted lines in different colors.

**b)** We used the same significance test method as **a)**, but we calculated the proportion of CTCF motifs involved in 15kb windows.

**Supplementary Tables:**

**Table S1. Statistics of genome assembly**
**Table S2. GO enrichment for lineage-specific orthologs**
**Table S3. Ohnolog group gene list**
**Table S4. number of amphixous orthologous groups with and without vertebrate ohnologs**
**Table S5. GO enrichment for genes in amphixous segmental duplications**
**Table S6. Sequencing data produced in this study**
**Table S7. The list of Bf fully sex-linked genes and their sex-biased expression**
**Table S8. The list of Bj fully sex-linked genes and their sex-biased expression**
**Table S9. QC of HiC data**
